# Lack of Consensus for Manual Mouse Sleep Scoring Limits Implementation of Automatic Deep Learning Models

**DOI:** 10.64898/2026.03.27.714381

**Authors:** Laura Rose, Alexander Neergaard Zahid, Javier García Ciudad, Christine Egebjerg, Louise Piilgaard, Frederikke Lynge Sørensen, Mie Andersen, Tessa Radovanovic, Anastasia Tsopanidou, Maiken Nedergaard, Sébastien Arthaud, Renato Maciel, Christelle Peyron, Chiara Berteotti, Viviana Lo Martire, Alessandro Silvani, Giovanna Zoccoli, Micaela Borsa, Antoine Adamantidis, Morten Mørup, Birgitte Rahbek Kornum

## Abstract

Scientists have for decades attempted to automate the manual sleep staging problem not only for human polysomnography data but also for rodent data. No model has, however, succeeded in fully replacing the manual procedure across clinics and laboratories. We hypothesize that this is due to the models’ limited ability to generalize to data from unseen laboratories. Our findings show that despite the high performance of four state-of-the-art models reported in initial publications, the published models struggle to generalize to other laboratories. We further show a significant improvement in model performance across labs by re-training them on a diverse dataset from five different sites. To assess the contribution of variability in manual scoring, ten experts from five laboratories all labelled the same nine mouse sleep recordings. The result revealed substantial scoring variability, particularly for rapid eye movement (REM) sleep, both within and between labs. In conclusion our study demonstrates that key challenges in the generalizability of state-of-the-art sleep scoring models are signal variability and label noise. Our study highlights the need for a standardized set of mouse sleep scoring guidelines to enable consistency and collaboration across the field. Until such a consensus is reached, we present four sufficiently robust models trained on diverse datasets that can serve as standardized tools across labs.

## Introduction

Mouse models are commonly used to study sleep due to the availability of tools that allow for genetic manipulation and sophisticated phenotyping. Like in humans, electroencephalogram (EEG) and electromyography (EMG) signals are used for classification of different vigilance states, and in mice these are typically divided into three stages: Wakefulness, NREM-sleep (NREMS), and REM-sleep (REMS). Although common sleep stage characteristics of mouse sleep, such as high delta power (0.5-4 Hz) in NREMS and high theta power (6-10 Hz) in REMS, are well established, inter-scorer variability still exists [1–3]. Unlike human sleep scoring, which follows the standardized guidelines of the American Academy of Sleep Medicine [4], there is no firm consensus for classifying wake and sleep stages in rodents [2].

Manual sleep stage classification in mice is still the gold standard for sleep research, despite it being highly time-consuming. This has prompted many scientists to explore automation of sleep stage classification in mice. Using automatic sleep scoring algorithms would not only accelerate sleep study evaluation but more importantly streamline the process across labs by providing a standardized tool for sleep scoring, enabling direct comparisons of results across different laboratories.

Variability in the EEG and EMG sleep signals presents a major challenge for automatic sleep stage classification. This variability arises both from biological and technical variation but likely also from label noise introduced by experts scoring sleep differently (subject bias). Technical variability reflects the wide range of experimental conditions across laboratories, as each lab typically uses unique hardware and recording setups (e.g., different number of electrodes, varying locations of the electrodes and different references). Even within a single lab, experiments vary due to biological factors in rodents such as genetic background, disease models, sex, and age. The label noise adds another layer of complexity, as a model will learn the specific scoring patterns from the training data, which limits its generalizability across datasets.

Researchers have applied various strategies to address this variability. Miladinovic et al. published the first deep learning model, SPINDLE, for mouse sleep staging in 2019 [1]. They argued that using a deep learning model would be a better way to approach the high variability rather than relying on handcrafted feature extraction and application of simple machine learning models as the high signal variability makes it challenging to define robust features that generalize well. They successfully achieved a generalization agreement of 97.39%, 93.21% and 97.47% in data cohorts of mutant mice generated from the same lab, of mice from an external lab, and of rats from an external lab, respectively. Later, Barger et al., introduced the mixture z-scoring as a part of their sleep stage classification model (SS-ANN) [5]. This approach aimed to preserve variability arising from experimental manipulations but corrected for the variability that arises from the signal acquisition [5]. As opposed to Miladinovic et al., Barger et al. aimed to reduce the learnable parameters from 6.8M parameters as in SPINDLE to 20K parameters in SS-ANN while maintaining generalizability. However, although they reported a held-out performance of 95-98%, the held-out data partly consisted of recordings from mice that also were a part of the training, which most likely inflated the performance.

Both SPINDLE and SS-ANN treat the sleep stage classification problem as an image classification task and use spectrograms for training the model. A different approach is to train a model on time-series EEG and EMG signals. This was applied in two other models generated by Grieger et al. [3] and Jha et al. [6]. Griger et al. further used data augmentation (i.e., through amplitude variability, frequency variability, EEG montage variability and time shift variability) to make their model robust to variations arising from data acquisition, scoring, and other unrelated physiological processes [3]. However, Grieger et al. found that they achieved better performance by omitting the data augmentation when they tested the model on a held-out in-house dataset. Jha et al. also used data augmentation in their model, SlumberNet based on the known ResNet model, and further attempted to obtain a robust classifier, that could work well across various sleep conditions by training a model on mice undergoing baseline sleep, sleep deprivation, and recovery sleep [6]. They showed that the model could perform well on an in-house mouse dataset across similar conditions. However, the model was not evaluated on external data cohorts.

As all four models originally were trained on data from a single laboratory which could limit their generalizability to other labs. Although models trained and evaluated on the same in-house datasets often achieve higher performance, this comes at the cost of reduced generalizability. Additionally, it artificially raises the performance benchmark for new publications. In contrast, models trained on diverse datasets are more resistant to overfitting to a single lab. While this may lead to slightly lower performance on in-house data, it is more likely to improve generalizability to external datasets and ultimately provide a more valuable tool for the field.

While much focus has been on signal variability, relatively little attention has been given to label noise stemming from variations in the manual labelling. Label noise introduces a ceiling effect on the performance of automatic sleep stage classification models, making it a crucial factor to address when developing robust and accurate sleep stage classification models. To address this, we collected a unique dataset where 10 experts (two experts from each lab) scored the same nine recordings, allowing us to examine the within- and between-lab variability. Our results highlight the need for a standardized set of mouse sleep scoring guidelines.

In conclusion we here demonstrate that training on diverse data from multiple labs and standardizing sleep scoring criteria across laboratories, rather than developing new models, is a critical first step towards fully automating sleep stage classification for mice.

## Results

### State-of-the-art sleep stage classification models do not generalize well when tested on external datasets

We first investigated to what extent state-of-the-art models generalized to unseen data from different laboratories. For this we chose the four state-of-the-art models described above (i.e., SPINDLE [1][5], SS-ANN [5], Grieger [3] and SlumberNet [6]). These four baseline models were selected based on the three following criteria: 1) it is a deep learning-based model, 2) the input to the model uses at least one EEG channel, and 3) the original data is available online (allowing for reproduction of the model). Further, data from five different laboratories (Cohort A – E) were collected resulting in a total of 83 wild-type (WT) mice with some mice having multiple recordings. Cohort A-E has previously been described in [12].

Each model was tested on cohort A-E to evaluate the robustness for classifying sleep stages on data from various labs (Figure 1). Our results show that both SPINDLE (Wakefulness -Recall: 98.8%, 93.7%, 97.1%, 96.7%, 95.1% across labs) and SS-ANN (Wakefulness - Recall: 90.9%, 93.0%, 96.2%, 92.0%, 96.9% across labs) perform consistently well in detecting Wakefulness across laboratories compared to Grieger (Wakefulness – Recall: 56.4%, 95.8%, 2.6%, 83.7%, 88.1%) and SlumberNet (Wakefulness – Recall: 94.3%, 60.3%, 83.7%, 36.7%, 12.12%). Both SPINDLE, SS-ANN and SlumberNet struggle in detecting REMS (REMS recall SPINDLE: 84.1%, 60.9%, 55.8%, 66.9%, 50.3%; REMS recall SS-ANN: 76.9%, 62.2%, 33.4% 51.1% and 77.5%; REMS recall SlumberNet 54.3%, 82%, 32.2%, 55.4%, 54.2%). The Grieger model achieves a recall for REMS above 90% for 4/5 labs (Recall REMS: 92.0%, 66.0%, 94.8%, 96%, 92.2%). A common trend across all models was that the sleep stage with the highest recall also was the most likely stage to be classified, suggesting that the high recall likely resulted from a tendency to classify the majority of the epochs into that particular stage. Our analysis shows that state-of-the-art models that have been trained on data from a single lab display highly variable performance on data from other laboratories.

**Figure 1.**
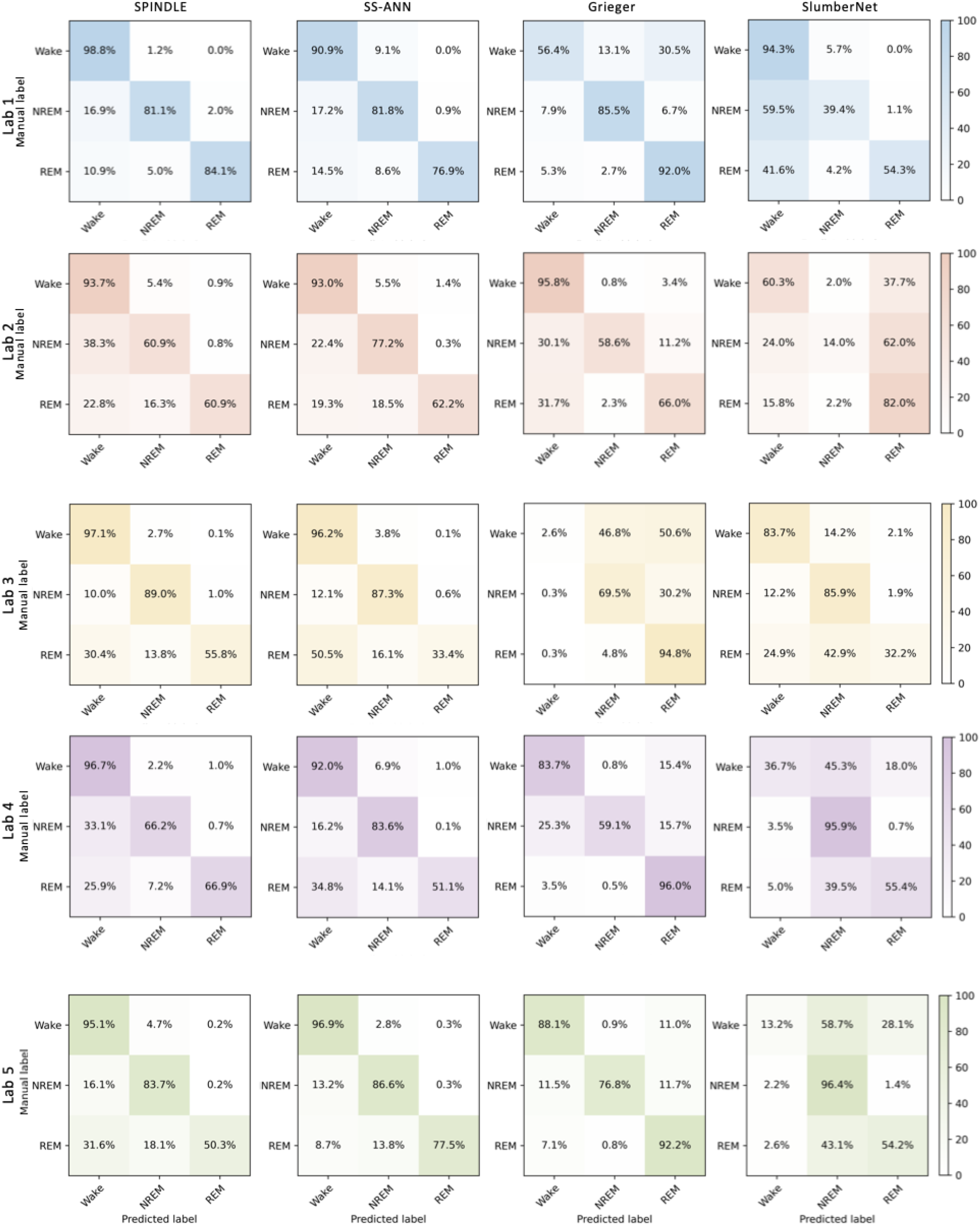
State-of-the-art-models do not generalize to external dataset. Confusion matrices of different state-of-the-art models tested across different laboratories (Lab 1-5) with row-wise normalization (recall). X-axis reflects the predictions and y-axis the manual labels. SPINDLE and SS-ANN are both trained on images, while Grieger and SlumberNet are trained on sequences of time-series signals.

### Models trained on a diverse dataset significantly improve performance when applied to unseen data

We aimed to address the generalization limitation of the state-of-the-art models by introducing a diverse training set from multiple laboratories, allowing the models to learn a broader range of scoring rules. We performed this in a leave-one-lab-out (LOLO) approach such that the models were trained on data from four labs and tested on the fifth lab in a cross-over fashion.

To evaluate the effect of this approach, we compared the results of three experiments: 1) performance of the *baseline* models trained on the original data and tested on cohort A-E; 2) performance of *fixed n* models trained on diverse data (identical in size to original data) from four different labs and tested on data from the fifth lab; and 3) performance of *all n* models trained on all data from four labs and tested on data from the fifth lab (Figure 2).

**Figure 2.**
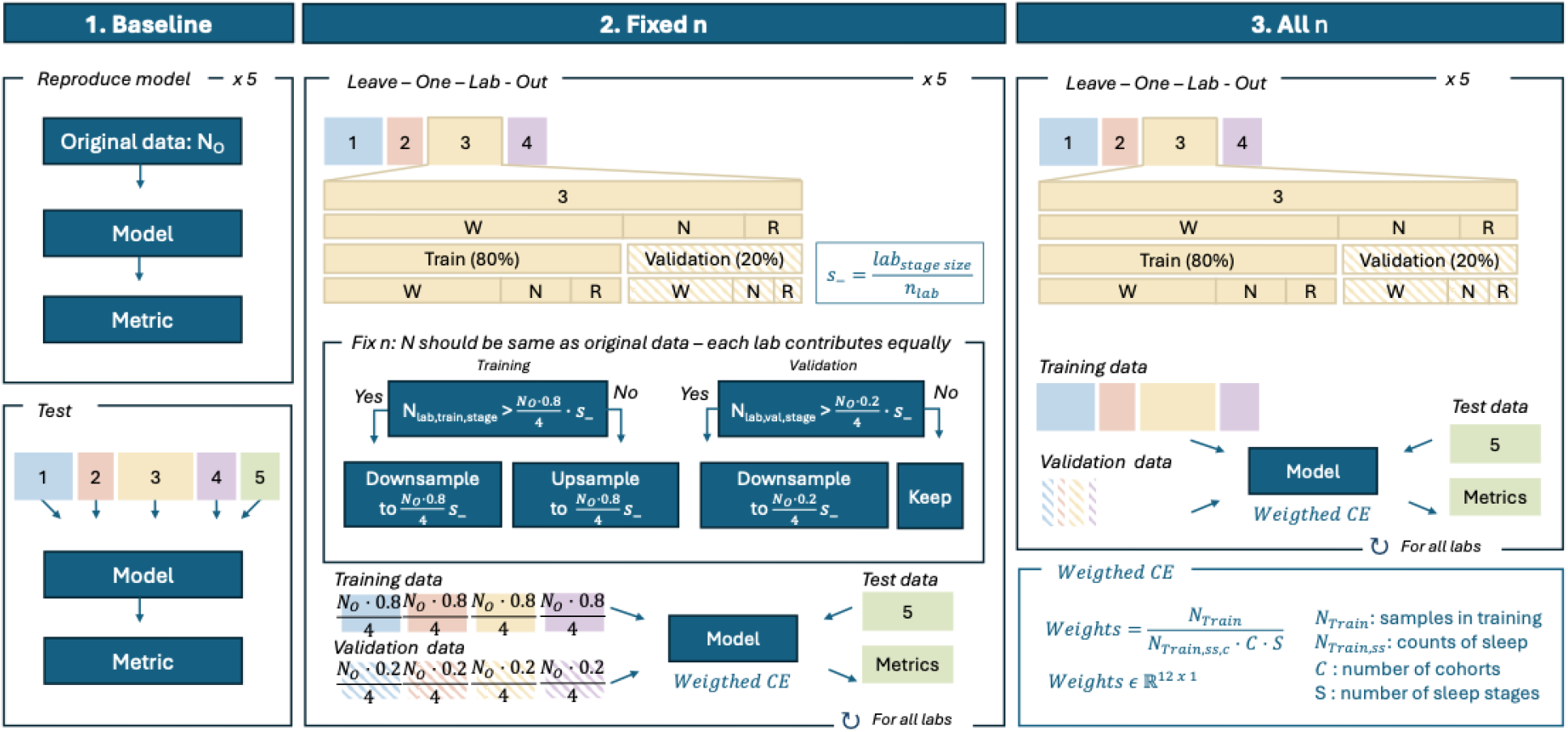
A flowchart illustrating three main experiments aimed at evaluating model performance across various scenarios: Testing whether state-of-the-art models can generalize to external datasets (Baseline). Assessing the improvement in model performance when trained on a diverse (Fixed n) or a large *and* diverse dataset (All n).

To test the effect of a diverse and large sample, we used a linear model to correct for confounding factors: lab and model. Our results demonstrate that training the models on a diverse dataset with a fixed sample size significantly improves overall performance when models were tested on unseen data (Figure 3; p-value 1.59 · 10^-6^). Similarly, training on a large, diverse sample also showed improved performance (Figure 3; p-value 1.07 · 10^-6^), with weight initialization accounted for in both cases. Although training on the diverse and large sample slightly improved model performance more than the diverse and fixed sample (Figure 3B; mean difference 0.0562 vs. 0.0572), the improvement between the fixed n (experiment 2) and all n (experiment 3) was not statistically significant. This demonstrates that the diversity of the training data matters more than the size of the training set when the goal is to achieve high generalizability of the model.

**Figure 3.**
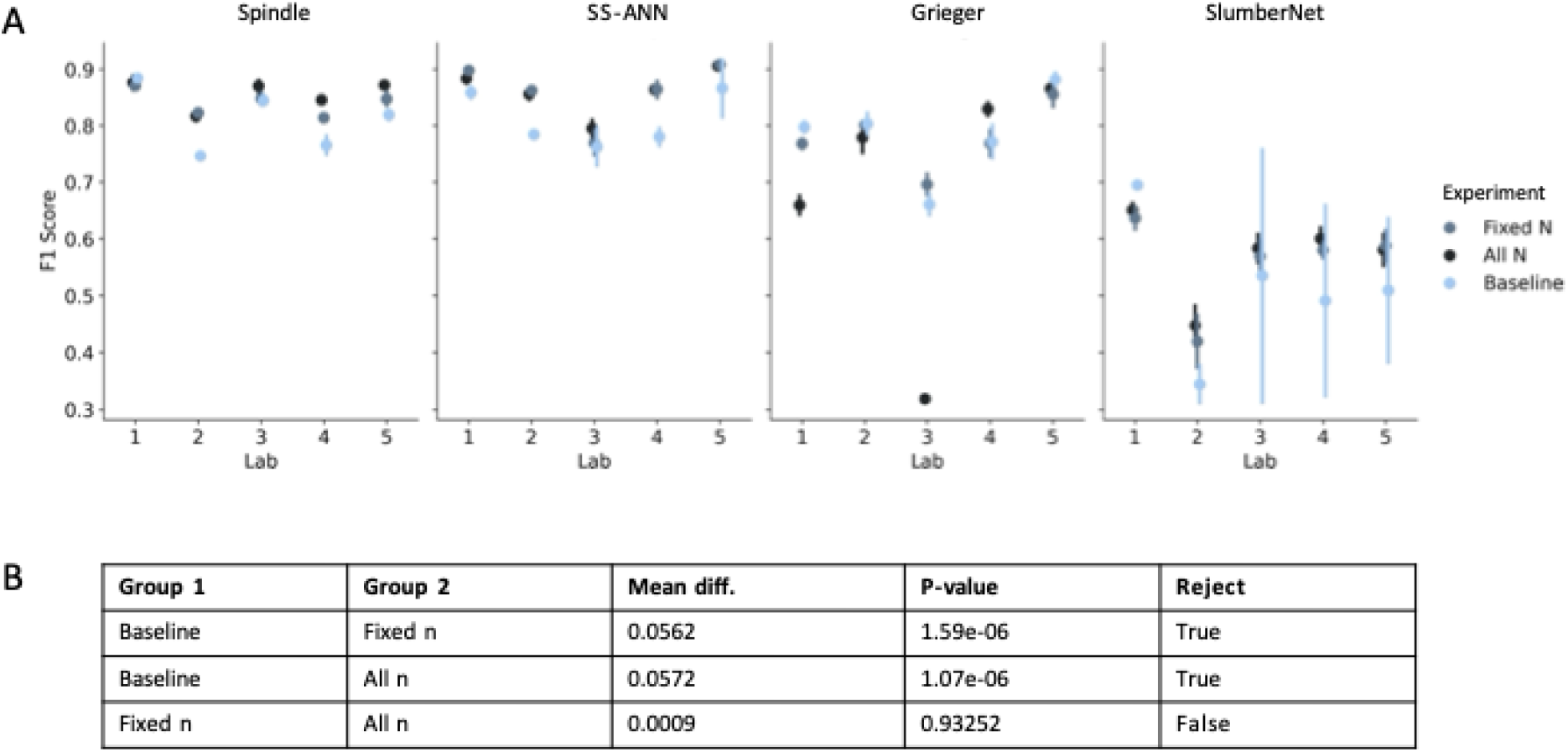
Having a diverse training sample significantly improve the state-of-the-art classifiers. (A) The macro F1-score (i.e., mean across sleep stages) for each experiment, lab and model. One sample represents the average macro F1-score across five repeated training runs and the error-bar reflects the standard deviation. (B) Mean differences between experimental groups, along with p-values and whether the null hypothesis could be rejected.

### Lack of consensus in manual sleep scoring within and between laboratories particularly for REMS

After observing the lack of generalizability of the state-of-the-art sleep scoring models, we decided to assess the label noise in manual scoring. To do so, we asked 10 experts, two from each lab, to manually annotate the same nine recordings. An example of an annotated signal is shown in Figure 4A The within-lab agreement was high for Wakefulness (Figure 4B; Cohen’s kappa = 0.93, 0.95, 0.91, 0.94, 0.91) compared to NREMS that was slightly lower (Figure 4B; Cohen’s kappa = 0.88, 0.94, 0.91, 0.91, 0.86), and to REMS (Figure 4B; Cohen’s kappa = 0.78, 0.93, 0.89, 0.90, 0.80), which showed the lowest agreement. The between-lab (i.e., experts from one lab were compared to experts from all other labs) agreement showed a similar pattern to the within-lab agreements (Figure 4C; Wakefulness, Cohen’s kappa = 0.90, 0.92, 0.91, 0.91, 0.90; NREMS, Cohen’s kappa = 0.87, 0.89, 0.89, 0.89, 0.87; REMS, Cohen’s kappa = 0.77, 0.87, 0.85, 0.86, 0.82). While the between-lab agreement was slightly lower than the within-lab agreement in all stages, the within-lab agreement highlights inconsistencies in scoring even between experts coming from the same lab.

**Figure 4.**
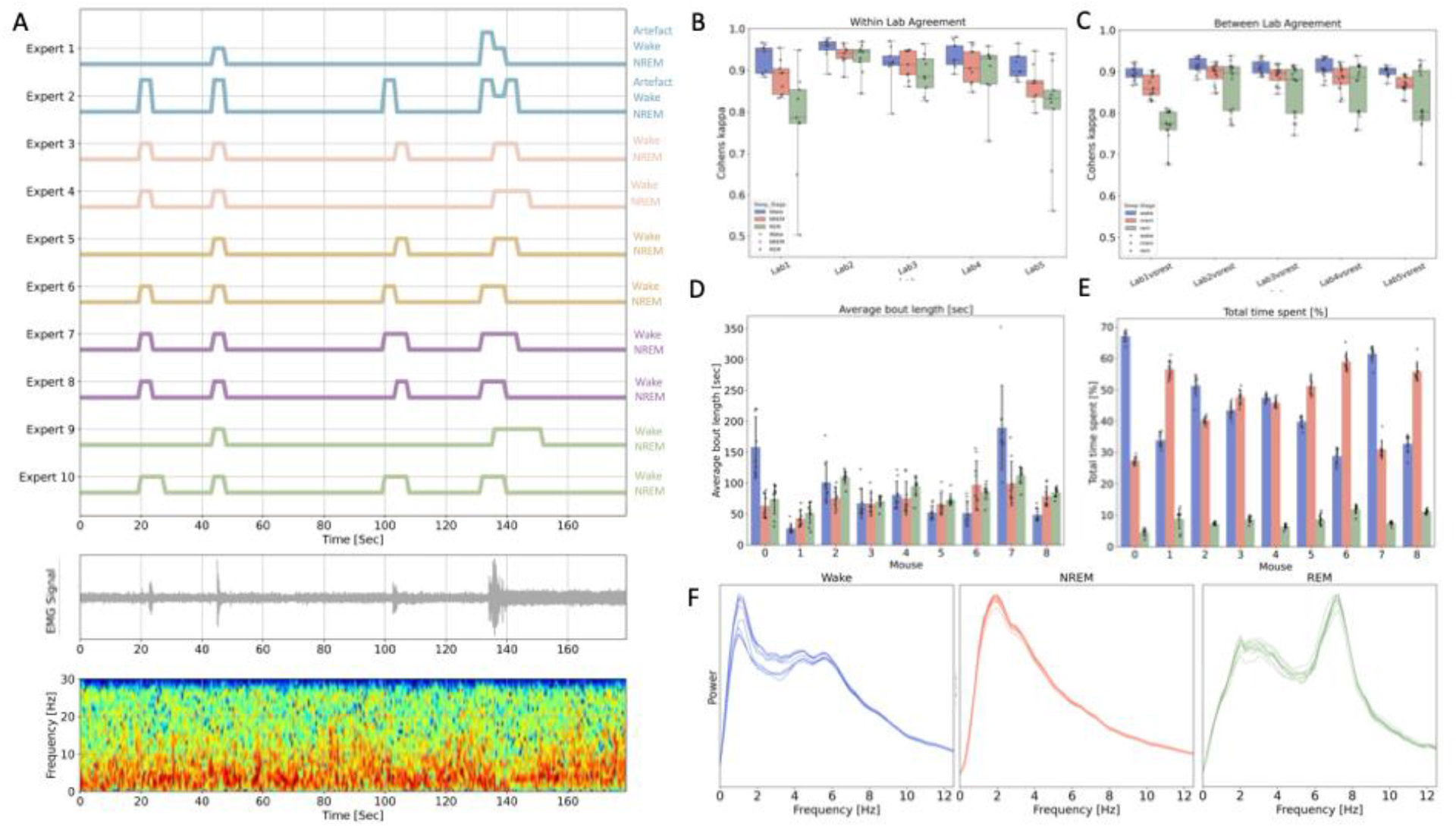
Lack of consensus in manual sleep scoring, both within and between labs, particularly for REM sleep. (A) Hypnograms from 10 experts of a three-minute EEG/EMG sequence. (B) Within lab agreement (i.e., Cohen’s kappa was calculated between two experts from the same lab) for each sleep stage and each mouse resulting in nine samples per sleep stage. (C) Between lab agreement (i.e., experts from one lab were compared to all other experts from other labs; 16 samples) for each sleep stage. Each sample reflects an average across recordings from five mice. Down-stream sleep analysis based on the manual labels from each of the 10 experts: (D) Average bout length [sec] and (E) Total time spent in each state [%]. Each point in graph D and E represents one expert. (F) Power spectrums of the EEG for each sleep stage. Each line represents one expert.

We further investigated how much the scoring variability affected downstream sleep analysis. Bout length analysis revealed that the average bout length varied more between experts than did the total time spent in each sleep stage (Figure 4D-E), suggesting that much of the variability occurs at the start and end of each bout, which is supported by visual inspection of the hypnogram (example in Figure 4A). Variability was also found within delta power during Wakefulness and within delta/theta power during REMS (Figure 4F).

### Manual label-derived hypnodensities show comparable macro sleep characteristics to those produced by the four fully trained deep learning models

After training the four models on all data from the five labs, we tested the models on data consisting of the nine recordings manually annotated by the 10 experts. We found that the macro sleep structure based on the hypnodensities from the manual sleep scores and the four sleep scoring models was similar across most models in both the dark and light phases (Figure 5; Figure S1-S4). Certain areas of the manual hypnodensity indicate a clear wake stage, while the deep learning models suggested brief wake-sleep mixed stages, which could be explained by the models being more likely to predict sleep stages (i.e., NREMS and REMS) compared to Wakefulness. This was further supported by calibration curves (Figure 5B: SPINDLE, SS-ANN, and Grieger), which showed that the models correctly predict wakefulness when the manual experts also reached a relatively high agreement on Wakefulness. In contrast, only a few experts had to agree on NREMS or REMS before the models predicted these stages with high certainty (Figure 5B: SPINDLE, SS-ANN, and Grieger). The calibration curve for the SlumberNet model indicated that this model, in contrast to the other three, was rather conservative in predicting both NREM and REMS, despite high agreement among experts.

**Figure 5.**
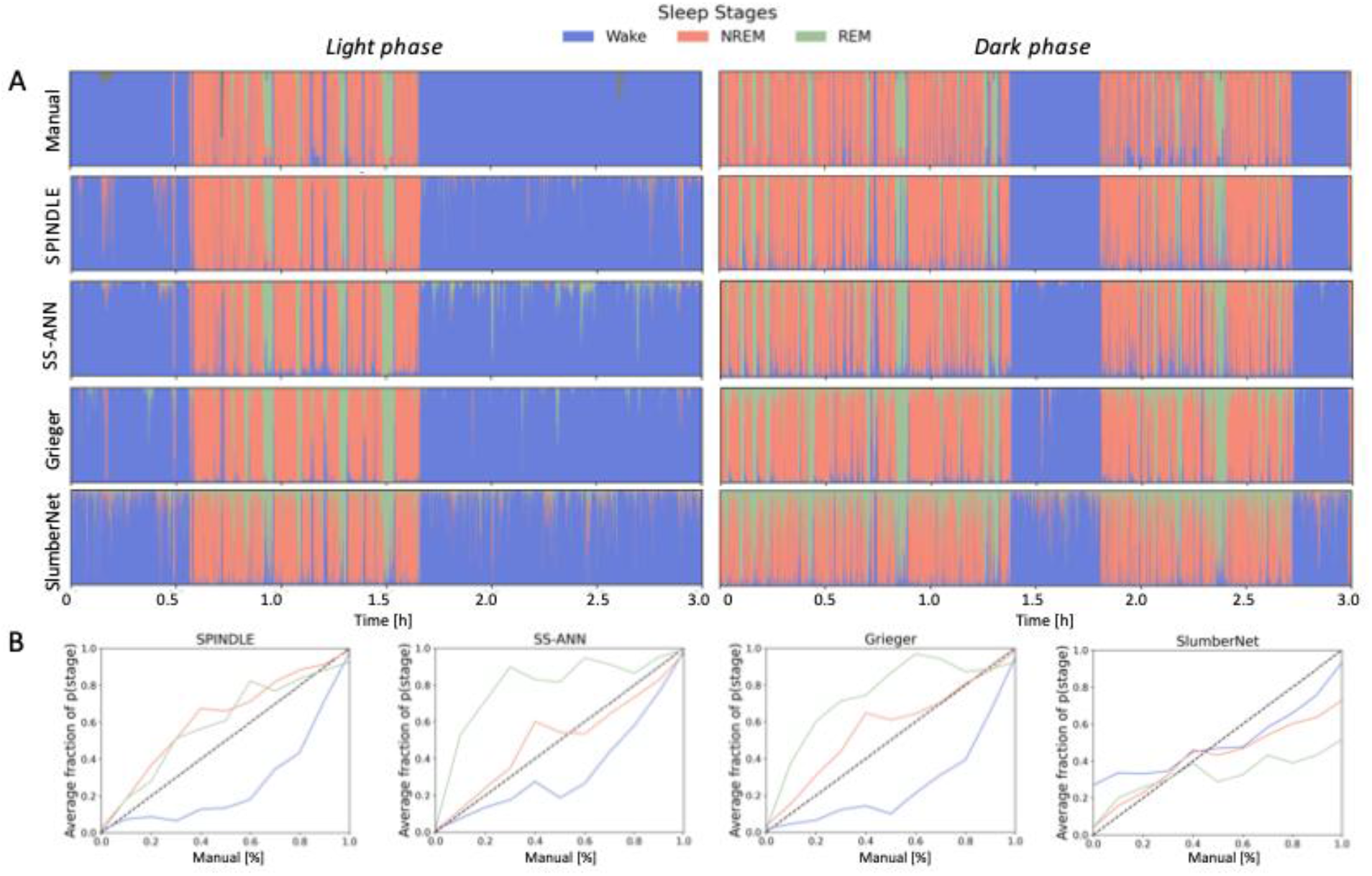
Hypnodensities derived from manual labels exhibit similar macro sleep characteristics to those generated by three of the four fully trained deep learning models. (A) The manual hypnodensities were derived based on the ten experts’ annotations for each mouse during dark phase and light phase. Hypnodensities based on the probability outputs from the fully trained SPINDLE, SS-ANN, Grieger and SlumberNet model. (B) Calibration curves for SPINDLE, SS-ANN, Grieger and SlumberNet. The x-axis reflects an aggregation of the 10 sets of manual labels (e.g., 2/10 experts agreed on NREM for a given epoch, 8/10 agreed on Wake, resulting in a probability distribution of p(Wake) = 0.8, p(NREM) = 0.2, and p(REM) = 0) labelled as manual [%]. The y-axis reflects the average fraction of epochs belonging to the sleep stage out of all epochs in the corresponding x-label bin. The black dashed line shows the optimal calibration curve.

## Discussion

In this study, we show that state-of-the-art sleep scoring models (when trained on data from a single lab) fail to generalize to data from other labs. However, we demonstrate that retraining these models on a diverse dataset significantly enhances their performance on unseen data. Finally, we highlight a lack of consensus in manual sleep scoring, particularly for REMS.

Although several automatic sleep stage classification models report high performance: SPINDLE with 99±0.5% accuracy [1], SS-ANN at 96.8% [5], Grieger with an F1-score of 0.95 [3], and SlumberNet at 97% accuracy [6], none of them perform consistently well across external laboratories (Figure 1). This inconsistency could come from signal variability or label noise. Furthermore, both SS-ANN and SlumberNet face the issue of information leakage (i.e., training and testing on the same mice), which can compromise the model’s generalizability and artificially inflate performance metrics.

When comparing the four models, SS-ANN is the only model that requires a few manually labelled epochs to calculate a standardized z-score, making it less flexible compared to SPINDLE and Grieger. Unlike the other models, SlumberNet does not include neighbouring epochs in its sequences, preventing it from leveraging contextual information. This absence of context could be one reason for its lower performance compared to the others, which use sequences of 20, 44, and 30 seconds for SPINDLE, SS-ANN, and Grieger, respectively. SPINDLE may offer more explainability, as it uses spectrograms instead of raw signal sequences as input. This approach allows for easier post-hoc explainability techniques to identify sleep markers and understand the rationale behind model predictions.

This trend of inflated performance has substantial implications for the field, as new models are typically expected to outperform established ones. Instead of focusing solely on performance within a single dataset, it may be more meaningful to evaluate model robustness across datasets, providing a better indication of real-world applicability. Another issue is that studies often report accuracy as the primary performance metric, which may not adequately capture generalizability. A high accuracy can be achieved by overclassifying the majority class, obscuring the actual model performance. Metrics such as the F1-score or balanced accuracy could provide a more accurate picture.

In this study, we re-trained four state-of-the-art models on data from five different laboratories, with the goal of creating more robust classifiers. By training all models on the same data, we revealed differences in how they classify sleep. For SPINDLE, SS-ANN, and Grieger, a strong agreement across expert labels corresponded with high prediction certainty from the models. In contrast, SlumberNet struggled to achieve high certainty, as reflected in the hypnodensities, which show more mixed-stage predictions compared to the other models (Figure 5). SPINDLE, SS-ANN, and Grieger tended to underestimate Wakefulness and overestimate sleep stages, potentially due to how the weighted loss function penalizes less frequent sleep stages more heavily, forcing the models to learn NREMS and REMS. Our results suggest that depending on the purpose of a given study, different models offer different advantages. An ensemble approach utilizing several models could also be considered, as the calibration curves indicate that some models overestimate NREMS while others overestimate REMS. Combining models may lead to more robust predictions and improve overall performance.

Our study raises the question of how we can expect the automatic sleep scoring models to generalize well, when manual labelling adds label noise to the data? We here show that training the models on a diverse dataset significantly improves the state-of-the-art model performance (Figure 3). However, diversity in the dataset (i.e., data from different labs) also means incorporating different sets of scoring rules, which may confuse the model. For example, one lab might classify most NREMS to REMS sleep transition epochs as NREMS, while another lab may label them as REMS. Thus, the lack of consensus for manual sleep scoring (Figure 4B-C), makes it difficult to develop a standardized tool that will be applicable to every laboratory.

EEG and EMG signals alone hold a considerable amount of variability, making them inherently challenging for models to learn. Adding label noise and lab-to-lab differences in data acquisition only makes the sleep staging problem more difficult. Although several labs report similar scoring criteria (Supplementary Table 1), disagreements persist (Figure 4B-C). This discrepancy may arise, in part, because experts not only rely on the signals but also frequently use the video of the mouse during scoring. In this study we focused on manual and automatic sleep stage classification of the three main sleep stages (i.e., Wakefulness, NREMS and REMS). Artifacts is another class that is handled very differently across laboratories. To advance the field, a discussion is needed on how to handle epochs where experts deviate from standard scoring criteria. Thus, the initial steps in automating sleep scoring may lie not in model development but in standardizing scoring criteria and hardware setups across laboratories to reduce these sources of variability, particularly during REMS.

In manual sleep staging, each epoch is assigned a discrete sleep stage (Wakefulness, NREMS, or REMS for mice). However, if a sleep stage is a mix of two stages, such as during transitions or in disease models like narcolepsy, it is not possible to represent this mixture without introducing new classes such as intermediate classes. With automatic sleep stage classification models, each epoch can be assigned a probability of belonging to multiple stages. However, model uncertainty is embedded within these probabilities, making it challenging to distinguish model uncertainty from biologically mixed stages. The graphical representation of sleep stage probabilities is called a hypnodensity [7]. This visual depiction of model uncertainties and mixed stages, which should become standard in the sleep field, is only possible with either a mix of labels from many experts or with automatic models, highlighting the additional information gained from using automatic sleep stage classification models.

Because we collected manual scores from 10 experts, we can examine whether areas of high uncertainty in the manual hypnodensity align with the mixed stages predicted by the model. Such mixed stages could be due to technical noise but could also represent biological meaningful mixed sleep stages. Future experiments should explore ways to distinguish model noise from biological mixed stages. This would potentially allow us to derive biomarkers from the hypnodensities in disease models.

## Methods

### Data collection overview

The work presented here is divided into three parts, each answering a different question:

1. Validation study: How well does the state-of-the-art sleep staging models generalize to unseen data?
2. Optimization study: To what extent can we improve the generalizability of the state-of-the-art models by training on a (larger and) more diverse dataset?
3. Consensus study: How consistent are the manual scores among experts within and between labs?

For the first two parts, data from five different laboratories (Cohort A-E) were collected resulting in a total of 83 wild-type (WT) mice with some mice having multiple recordings. For the third study, two EEG/EMG recordings were collected from five WT mice and scored by 10 experts from five different laboratories. An overview of the data is presented in Table 1.

**Table 1.** Data overview.

| Cohort | Number of mice | EEG | EMG | Number of epochs | Scored by n experts |
| --- | --- | --- | --- | --- | --- |
| A - lab 1 | 10 | 1 | 1 | 619534 | 1 |
| B - lab 2 | 17 | 4 | 1 | 184500 | 1 |
| C - lab 3 | 23 | 2 | 1 | 885404 | 1 |
| D - lab 4 | 28 | 1 | 1 | 74546 | 1 |
| E - lab 5 | 5 | 2 | 1 | 194307 | 1 |
| F | 5 | 2 | 1 | 24300 | 10 |

Cohort A-E has previously been described [12], while Cohort F is unique to this paper and described below. All analyses were performed using either Python or MATLAB 2023b, depending on the programming language of the original model. The specific environments, including package versions, are detailed in the supplementary materials for each model (suppl. data S1-S3).

All four fully trained models have been made available online https://github.com/laulaurose.

### Data collected specifically for this study (Cohort F)

We acquired EEG/EMG recordings from five young adult female mice (nine weeks old, C57BL/6JTac mice). EEG/EMG surgery was performed as described in Egebjerg et al. [8] with a two-EEG and one-EMG channel head-mount (#8201; Pinnacle Tech. Inc., KS, USA). Three days before the recordings, the mice were habituated to a 10” circular transparent recording cage (#9000-K20; Pinnacle Tech. Inc., KS, USA) with a video camera attached outside the cage. Four of the five mice had two EEG/EMG/video recordings each (one during the light phase and one during the dark phase), each lasting three hours. Due to muscle artifacts disrupting the EEG signal, the light phase recording was excluded for the fifth mouse, resulting in a total of 9 recordings (five dark phase and four light phase) included for scoring.

This data was distributed to five laboratories, where two experts from each lab annotated the recordings, resulting in 10 independent scorings per recording. Sleep stages: Wakefulness, NREMS, and REMS were scored by the experts, following local scoring guidelines as summarized in Supplementary Table 1. The scoring guideline for our lab are presented in Table 2). Most laboratories scored standard sleep stages and artifacts, however, one laboratory also classified intermediate states (epochs showing characteristics of two states simultaneously).

**Table 2.**
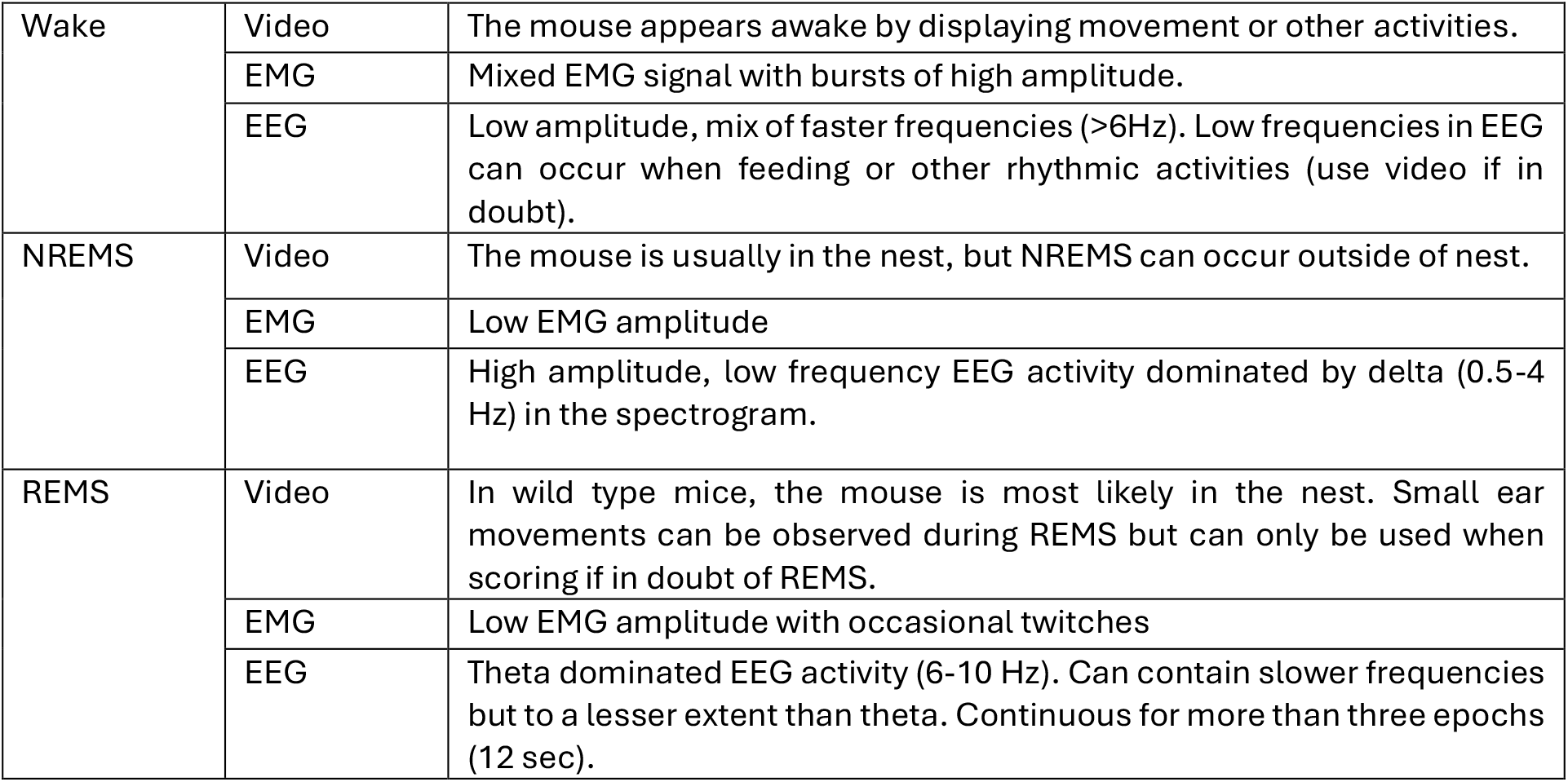
Scoring guideline for sleep stages from our lab.

|  |  |  |
| --- | --- | --- |
| Wake | Video | The mouse appears awake by displaying movement or other activities. |
|  | EMG | Mixed EMG signal with bursts of high amplitude. |
|  | EEG | Low amplitude, mix of faster frequencies (>6Hz). Low frequencies in EEG can occur when feeding or other rhythmic activities (use video if in doubt). |
| NREMS | Video | The mouse is usually in the nest, but NREMS can occur outside of nest. |
|  | EMG | Low EMG amplitude |
|  | EEG | High amplitude, low frequency EEG activity dominated by delta (0.5-4 Hz) in the spectrogram. |
| REMS | Video | In wild type mice, the mouse is most likely in the nest. Small ear movements can be observed during REMS but can only be used when scoring if in doubt of REMS. |
|  | EMG | Low EMG amplitude with occasional twitches |
|  | EEG | Theta dominated EEG activity (6-10 Hz). Can contain slower frequencies but to a lesser extent than theta. Continuous for more than three epochs (12 sec). |

### Reproduction of deep learning models and preprocessing pipelines

We reproduced four state-of-the-art deep learning models (i.e., baseline models; Figure 1) and tested the model performance in cohort A-E. The models are briefly explained along with the preprocessing pipeline below. We have used the same preprocessing pipeline and model architecture as reported in the original papers, unless otherwise specified.

Each baseline model was reproduced by training the model on its original dataset (Supplementary Table 2). To eliminate the effect of random weight initialization, the models were each reproduced five times using different random initializations. If the model weights were available online, we used it as the “baseline” model for testing. Otherwise, we selected the model that most closely matched the performance reported in the paper. These models were then used to test the performance on external data cohorts (cohort A-E). We chose the same hyperparameters (i.e., learning rate, batch size, and number of epochs) as originally reported in the papers.

### SPINDLE

The SPINDLE model [1] originally consists of two convolutional neural networks (CNN), one for sleep staging and one for artifact detection. The framework further involves an integration of a Hidden Markov Model (HMM) as a post-hoc analysis to the CNN for sleep staging. The HMM serves as a model that corrects for biological infeasible transitions such as Wakefulness to REMS. Since most laboratories did not score artifacts, training a second CNN for artifact detection was not feasible. Furthermore, as our goal is to evaluate the performance of sleep classification models without post hoc modifications that could artificially enhance results, only the sleep staging CNN was relevant to our study. Thus, moving forward, the SPINDLE model exclusively refers to the CNN classifier for sleep staging.

The SPINDLE model uses two EEG channels and one EMG channel. In cases where only one EEG channel was available, the same EEG would be used twice. Both signals were resampled to 128 Hz before applying the short-time Fourier transform to obtain spectrograms. The EEG spectrograms were bandpass filtered, log-scaled and standardized per frequency component. For the EMG spectrogram the PSD was summed within 0.5-30 Hz yielding a 1d array that was replicated to fit the dimension of the EEG spectrograms. The multi-channel EEG and EMG were concatenated and segmented into periods of five epochs, such that the epoch of interest was centred.

### SS-ANN

SS-ANN uses one EEG channel and one EMG channel. In cases where more than one EEG channel was available, random sampling was used to choose an electrode. Both signals were down-sampled to 128 Hz. The EEG signal was Fourier transformed to obtain spectrograms that were further down-sampled by a factor of two between 20-50 Hz for parameter reduction purposes. The EMG signal was bandpass filtered, and the root mean square (rms) was calculated for each epoch yielding a 1d array that was repeated nine times. The spectrogram (176 x n) was concatenated with the rms of the EMG signal (9 x n) resulting in 185 features per epoch, with n representing sequences of 9 epochs with the epoch of interest centred. The features were further standardized with the mixture z-score, as reported in [5].

### Grieger model

In contrast to other models, the Grieger model aims to classify additional stages such as pre-REM sleep and artifacts based on a single EEG channel [3]. Here, we refer to a modified version of the Grieger model, which only classifies the three main stages: Wakefulness, NREMS, and REMS. Additionally, we did not use the data augmentation approaches from the original paper as this was shown to decrease model performance. Finally, rather than artificially rebalancing the training set, we rebalanced the training data through a weighted loss function. The data was low-pass filtered with a cutoff frequency of 25.6 Hz, downsampled to 64 Hz, and z-scored per recording. The input to the model was a signal segment of 30 seconds, with the epoch of interest at the centre of the segment.

### SlumberNet

SlumberNet [6] uses one EEG and one EMG channel. The signals were first split into sequences of one epoch. Each epoch was later down-sampled to 64 Hz and the EMG signal was additionally high-pass filtered (cutoff frequency of 16 Hz) and further baseline-corrected as noted in [6]. A few modifications were added to the model. We identified a reshaping error in the original paper, where the input was incorrectly formed by concatenating two sets of 2-second EEG and 2-second EMG signals, instead of the intended 4-second EEG and 4-second EMG signals as described. We have since corrected this mistake and implemented the intended 4-second input format. Additionally, we added a standardization step to the preprocessing pipeline to address differences in signal acquisition across labs (e.g., variations in electrode placement and hardware), which can impact the signal amplitude. For each lab, we computed the overall mean and standard deviation of all epochs. We then standardized each epoch by subtracting the mean and dividing by the lab-specific standard deviation.

### Training the models on a diverse dataset

By leveraging data from five different labs, the models were trained using a leave-one-lab-out (LOLO) approach, resulting in five distinct models, with each left-out lab serving as the test set (Supplementary Table 2). We further investigated how much the models would benefit from being trained on a diverse and large dataset (i.e., all n; Figure 2). As a final step we trained all four sleep staging classifiers on all data and validated their performance using an external dataset (Cohort F).

In a fixed n experiment, the sample size was fixed to that of the original model, but the dataset was replaced with a more diverse dataset (cohorts A–E). To ensure that each lab contributed with an equal amount of data while keeping the original sample size *N*_*0*_, the following stratification was used. Data from each lab was first split into a training (i.e., 80%) and validation set (i.e., 20%; Figure 2). For the validation set: if 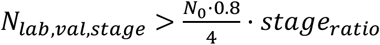 (Where 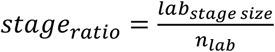) the data would be downsampled by random sampling of *N*_*lab*_,_*val*_,_*stage*_. For the training set: if 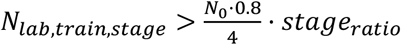, the data would be random sampled, else it would be upsampled. The models were trained with a weighted loss function to balance classes. As for the baseline models, the LOLO training was repeated five times to take different weight initializations into account.

Next, using the same LOLO framework, we allowed the models to utilize all available training data from each cohort. The data was split for each lab into a training and validation set in an 80%-20% ratio. We applied the same weighted loss function as in previous experiments. However, given the varying contributions of data from each lab, the weights further accounted for both imbalanced cohort sizes and differences in sleep stage distributions. Each model was trained five times per lab, resulting in a total of 25 models (i.e., five runs per model).

As a final step, we trained all four models (i.e., SPINDLE, SS-ANN, Grieger and SlumberNet) on all data (cohort A-E) without leaving any lab out, to obtain four final models.

### Addressing within and between lab variability

To assess the manual sleep scoring agreement both within and between labs, Cohen’s κ[9] was used as a metric to reflect the agreement both within and between labs. Cohen’s κ is based on true positives (TP), true negatives (TN), false negatives (FN) and false positives (FP):

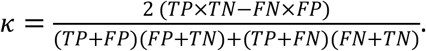

We calculated Cohen’s κ for each sleep stage and then averaged across stages for each mouse. For the within-lab analysis, two experts were compared for each mouse. To assess the between-lab agreement, we compared the agreement between lab i and the remaining four labs. Thus, for a single study, the sleep scoring from expert 1 from lab 1 was compared with all experts excluding expert 2 from lab 1. This was repeated for expert 2 from lab 1 resulting in 16 comparisons. We then computed the average agreement across all mice yielding 16 samples for comparison. To examine the effect of the sleep scoring variability on downstream analysis, we calculated features such as EEG power, average bout length, and the total time spent in each sleep stage for each set of sleep annotations. We used Welch’s method with a window size of 4 seconds (SciPy 1.7.3 [10]) to compute the power spectrums. For the bout analysis, the total time spent in each stage was calculated as the total time spent in that stage as a fraction of total time spent in all stages, and the average bout length was calculated in seconds.

### Comparing manual vs. automatic sleep stage classification

For each mouse, we converted the 10 sets of expert labels into a probability matrix, such that if 2/10 experts agreed on NREMS for a given epoch and 8/10 agreed on Wakefulness, the resulting probability distribution would be *p*(wakefulness) = 0.8, *p*(NREMs) = 0.2, and *p*(REM) = 0). We used these probability matrices, along with the probability outputs from the four fully trained deep learning models, to compute calibration curves for each model and each sleep stage. The calibration curves were derived by counting the number of epochs correctly classified as the actual sleep stage within that manual probability bin and dividing it by the total number of epochs in that bin.

### Statistics

We have used the macro F1-score to evaluate the performance of all automatic sleep stage classifiers, where

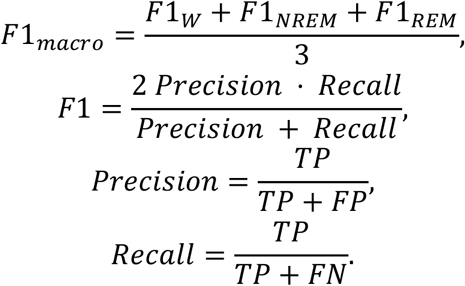

As we ran all three experiments five times for each model, we obtained 15 sets of predictions for each lab and model. To examine whether it helps the automatic sleep scoring models to learn a more diverse set of scoring rules, we tested the null hypotheses

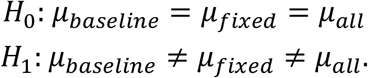

We fitted a linear model to account for confounding factors (i.e., lab and model),

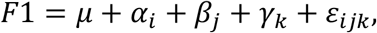

where *μ* represents the overall mean, *α*_*i*_ the effect of the model, *β*_*j*_ the effect of the lab, and *γ*_*k*_ the effect of the experiment. We used the OLS function from statsmodel [11] to fit the linear model. We also investigated a model with interaction terms, but did not observe any statistical difference.

## Supporting information

environment_grieger

environment_slumber

environment_spindle

## Data availability statement

The datasets generated and/or analysed during the current study are available in the Openneuro repository, https://openneuro.org/datasets/ds006366

## Supplementary figures and information Rose et al

### List of supplementary materials

**Figure S1.**
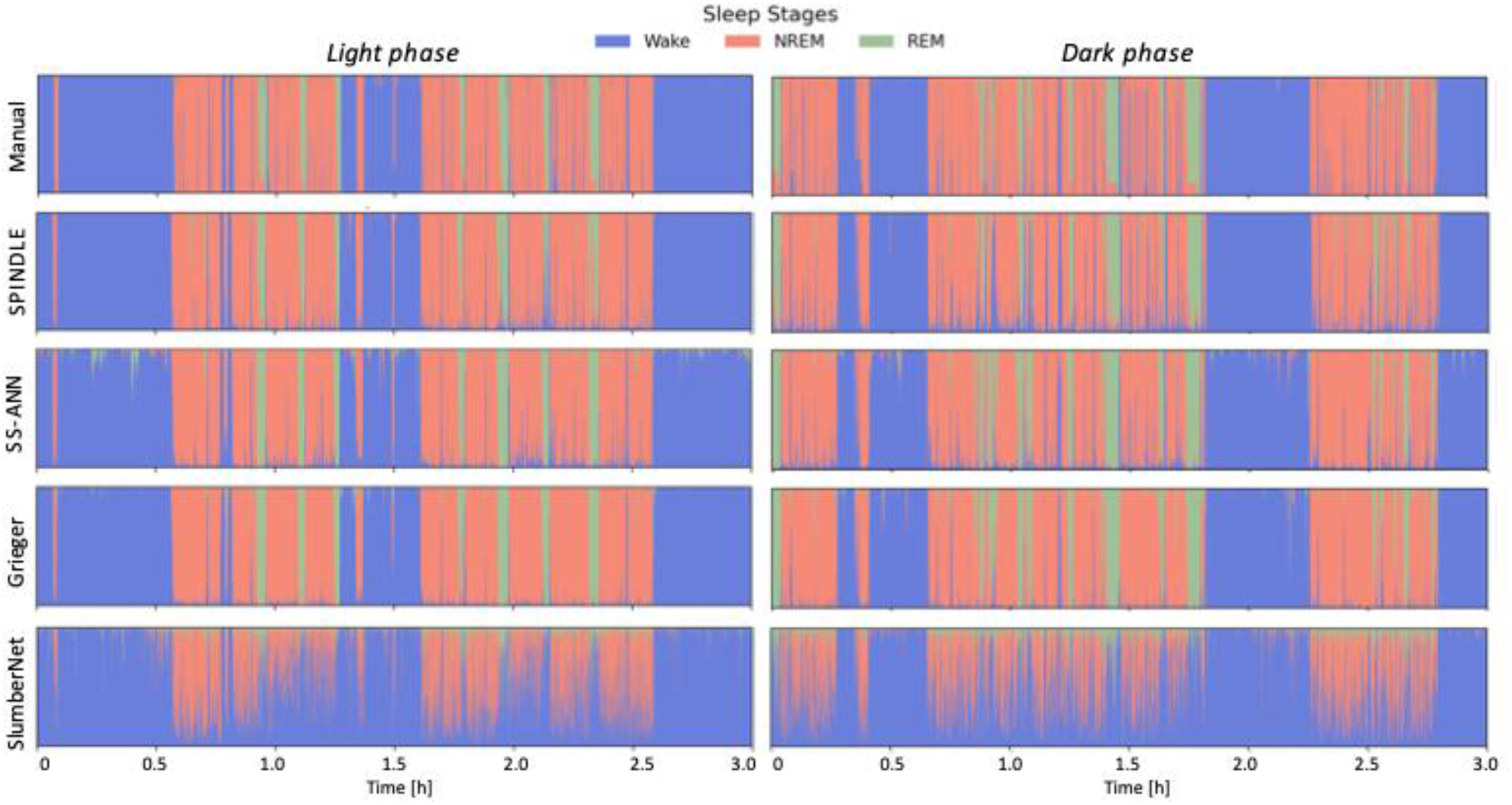
Hypnodensity plots from 2 x 3-hour recording of a single mouse, generated using manual labels and predictions from the four fully trained models.

**Figure S2.**
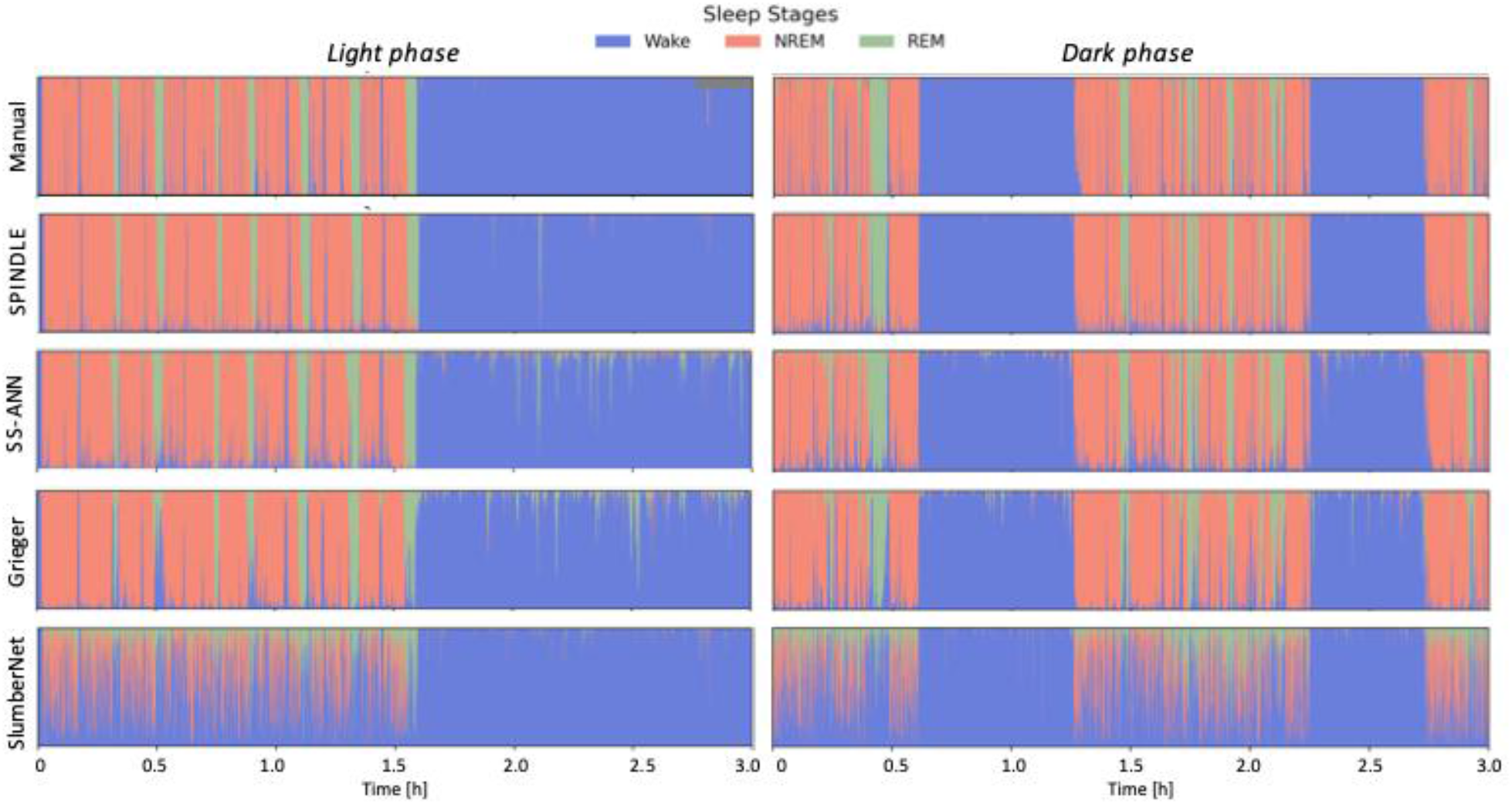
Hypnodensity plots from 2 x 3-hour recording of a single mouse, generated using manual labels and predictions from the four fully trained models.

**Figure S3.**
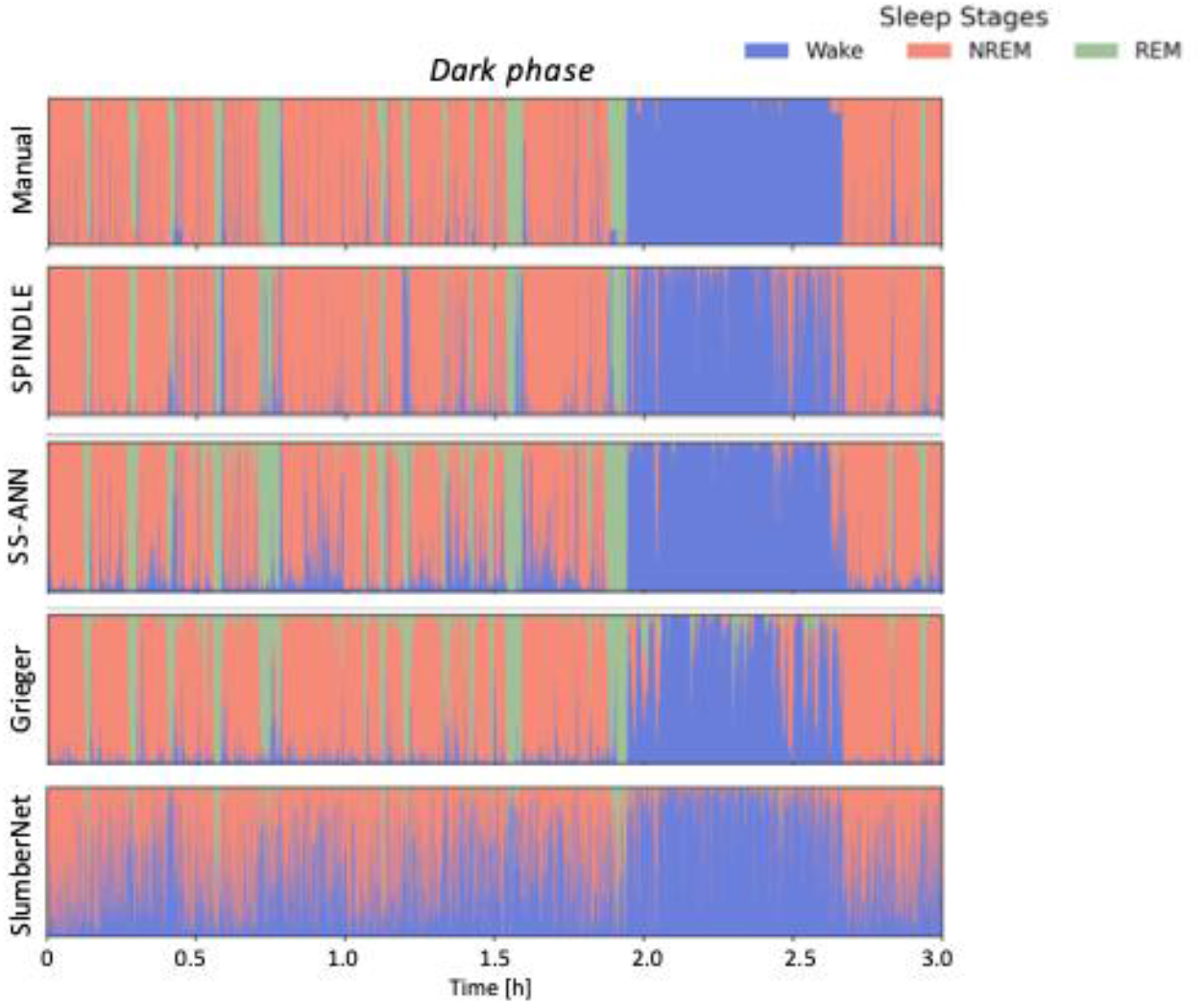
Hypnodensity plots from a 3-hour recording of a single mouse, generated using manual labels and predictions from the four fully trained models.

**Figure S4.**
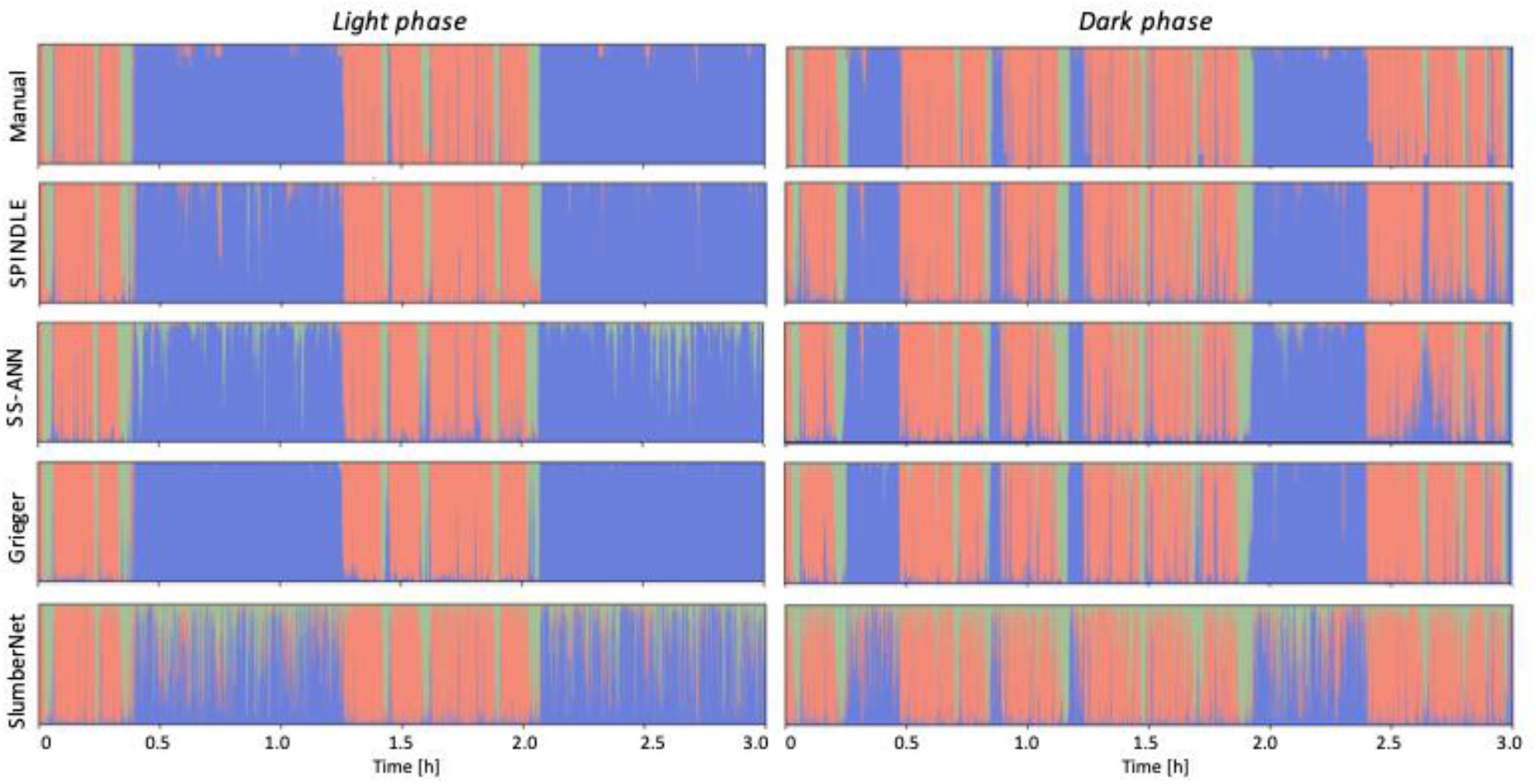
Hypnodensity plots from 2 x 3-hour recording of a single mouse, generated using manual labels and predictions from the four fully trained models

**Supplementary Table 1.** An overview of the scoring criteria of Wakefulness, NREMS, and REMS for each lab.

|  | Wakefulness |  | NREMS |  | REMS |  |
| --- | --- | --- | --- | --- | --- | --- |
|  | EEG | EMG | EEG | EMG | EEG | EMG |
| Cohort A | Low voltage with possible delta (0.5-4 Hz) and theta (6-9 Hz) frequency components | High EMG tone | EEG at high voltage with prominent delta (0.5-4 Hz) frequency components | EMG tone lower than in wakefulness | EEG at low voltage with predominant theta (6-9 Hz) frequency components | EMG indicating muscle atonia with occasional muscle twitches |
| Cohort B | Periods with low amplitude desynchronized EEG signals | High, tonic EMG activity containing phasic bursts | Synchronized, high amplitude oscillations in the slow (<1 Hz) and delta (1-4 Hz) band | Low EMG tone without bursts | Pronounced theta (5-10 Hz) oscillations or an increased amount of desynchronized and high-frequency oscillation | Almost complete absence of EMG tone except short twitches |
| Cohort C | Low voltage; mixed frequency $\theta$ (6-10 Hz), alpha (8-14Hz), beta (15-30Hz) and gamma (30-120Hz) | High amplitude | High amplitude low frequency EEG activity $\delta$ (0.5-4 Hz) | Low EMG amplitude | Low voltage fast EEG activity; theta dominated activity $\theta$ (6-10 Hz) | Low EMG amplitude with occasional muscle twitches |
| Cohort D | Low EEG amplitude. Low Fast Fourier Transform (FFT) magnitude across all frequency bands < 30 Hz | High EMG amplitude with phasic events | High amplitude with high FFT magnitude in delta range (0.5-4 Hz) relative to the other frequency bands | Low amplitude relative to that in wakefulness without phasic events | Low amplitude with relatively high FFT magnitude in theta range (4-8 Hz) compared to other bands | Low amplitude relative to wakefulness but with muscle twitches |
| Cohort E | Desynchronized (activated) low-amplitude EEG | Tonic and phasic EMG activity | High-voltage slow-waves (0.5-4Hz) | Loss of phasic EMG activity with low EMG tone | Regular and pronounced theta rhythm (6-9Hz) on parietal EEG | Absence of phasic and tonic EMG activity, except short twitches |

**Supplementary Table 2.** Overview of the different experiments that were conducted with specifications of the sample size and dataset that served as either training or test set. The table only lists experiments for one model (SPINDLE). The same strategy was used for all four models.

| Experiment | Model | Sample size | Train | Test | # of repeats |
| --- | --- | --- | --- | --- | --- |
| Baseline | SPINDLE | $N_{\text{original}}$ | Original data | Lab 1 | 5 |
| Baseline | SPINDLE | $N_{\text{original}}$ | Original data | Lab 2 | 5 |
| Baseline | SPINDLE | $N_{\text{original}}$ | Original data | Lab 3 | 5 |
| Baseline | SPINDLE | $N_{\text{original}}$ | Original data | Lab 4 | 5 |
| Baseline | SPINDLE | $N_{\text{original}}$ | Original data | Lab 5 | 5 |
| Fixed n | SPINDLE | $N_{\text{original}}$ | Lab2, lab3, lab4, lab5 | Lab 1 | 5 |
| Fixed n | SPINDLE | $N_{\text{original}}$ | Lab1, lab3, lab4, lab5 | Lab 2 | 5 |
| Fixed n | SPINDLE | $N_{\text{original}}$ | Lab1, lab2, lab4, lab5 | Lab 3 | 5 |
| Fixed n | SPINDLE | $N_{\text{original}}$ | Lab1, lab2, lab3, lab5 | Lab 4 | 5 |
| Fixed n | SPINDLE | $N_{\text{original}}$ | Lab1, lab2, lab3, lab4 | Lab 5 | 5 |
| All n | SPINDLE | $N_{\text{All}}$ | Lab2, lab3, lab4, lab5 | Lab 1 | 5 |
| All n | SPINDLE | $N_{\text{All}}$ | Lab1, lab3, lab4, lab5 | Lab 2 | 5 |
| All n | SPINDLE | $N_{\text{All}}$ | Lab1, lab2, lab4, lab5 | Lab 3 | 5 |
| All n | SPINDLE | $N_{\text{All}}$ | Lab1, lab2, lab3, lab5 | Lab 4 | 5 |
| All n | SPINDLE | $N_{\text{All}}$ | Lab1, lab2, lab3, lab4 | Lab 5 | 5 |
| Final model | SPINDLE | $N_{\text{Al}}$ | Lab1, lab2, lab3, lab4, lab5 | Cohort F | 1 |

## Further provided

**Environment.txt file for SPINDLE model**

**Environment.txt file for Griger model**

**Environment.txt file for SlumberNet model**

