## Supplementary material for "Lack of Consensus for Manual Mouse Sleep Scoring Limits Implementation of Automatic Deep Learning Models": environment_grieger

**Data S2: Environment.txt file for Grieger model**

name: grieger

channels:

- pytorch

- defaults

dependencies:

- _libgcc_mutex=0.1=main

- blas=1.0=mkl

- blosc=1.19.0=hd408876_0

- brotlipy=0.7.0=py37h27cfd23_1003

- bzip2=1.0.8=h7b6447c_0

- ca-certificates=2024.3.11=h06a4308_0

- certifi=2022.12.7=py37h06a4308_0

- cffi=1.15.0=py37hd667e15_1

- charset-normalizer=2.0.4=pyhd3eb1b0_0

- cryptography=39.0.1=py37h9ce1e76_0

- cudatoolkit=10.0.130=0

- cycler=0.10.0=py37_0

- dbus=1.13.16=hb2f20db_0

- expat=2.2.9=he6710b0_2

- ffmpeg=4.3=hf484d3e_0

- flit-core=3.6.0=pyhd3eb1b0_0

- fontconfig=2.13.0=h9420a91_0

- freetype=2.10.2=h5ab3b9f_0

- glib=2.65.0=h3eb4bd4_0

- gmp=6.2.1=h295c915_3

- gnutls=3.6.15=he1e5248_0

- gst-plugins-base=1.14.0=hbbd80ab_1

- gstreamer=1.14.0=hb31296c_0

- h5py=2.10.0=py37h7918eee_0

- hdf5=1.10.4=hb1b8bf9_0

- icu=58.2=he6710b0_3

- idna=3.4=py37h06a4308_0

- intel-openmp=2020.1=217

- joblib=0.15.1=py_0

- jpeg=9b=h024ee3a_2

- kiwisolver=1.2.0=py37hfd86e86_0

- lame=3.100=h7b6447c_0

- ld_impl_linux-64=2.33.1=h53a641e_7

- libedit=3.1.20191231=h7b6447c_0

- libffi=3.3=he6710b0_1

- libgcc-ng=9.1.0=hdf63c60_0

- libgfortran-ng=7.3.0=hdf63c60_0

- libiconv=1.16=h7f8727e_2

- libidn2=2.3.2=h7f8727e_0

- libpng=1.6.37=hbc83047_0

- libstdcxx-ng=9.1.0=hdf63c60_0

- libtasn1=4.16.0=h27cfd23_0

- libtiff=4.1.0=h2733197_1

- libunistring=0.9.10=h27cfd23_0

- libuuid=1.0.3=h1bed415_2

- libxcb=1.14=h7b6447c_0

- libxml2=2.9.10=he19cac6_1

- lz4-c=1.9.2=he6710b0_0

- lzo=2.10=h7b6447c_2

- matplotlib=3.2.2=0

- matplotlib-base=3.2.2=py37hef1b27d_0

- mkl=2020.1=217

- mkl-service=2.3.0=py37he904b0f_0

- mkl_fft=1.1.0=py37h23d657b_0

- mkl_random=1.1.1=py37h0573a6f_0

- mock=4.0.2=py_0

- ncurses=6.2=he6710b0_1

- nettle=3.7.3=hbbd107a_1

- ninja=1.9.0=py37hfd86e86_0

- numexpr=2.7.1=py37h423224d_0

- numpy=1.18.5=py37ha1c710e_0

- numpy-base=1.18.5=py37hde5b4d6_0

- olefile=0.46=py37_0

- openh264=2.1.1=h4ff587b_0

- openssl=1.1.1w=h7f8727e_0

- pandas=1.2.4=py37ha9443f7_0

- pcre=8.44=he6710b0_0

- pillow=7.1.2=py37hb39fc2d_0

- pip=20.1.1=py37_1

- pycparser=2.21=pyhd3eb1b0_0

- pyopenssl=23.0.0=py37h06a4308_0

- pyparsing=2.4.7=py_0

- pyqt=5.9.2=py37h05f1152_2

- pysocks=1.7.1=py37_1

- pytables=3.6.1=py37h71ec239_0

- python=3.7.7=hcff3b4d_5

- python-dateutil=2.8.1=py_0

- pytorch=1.13.1=py3.7_cpu_0

- pytorch-mutex=1.0=cpu

- pytz=2022.7=py37h06a4308_0

- pyyaml=5.3.1=py37h7b6447c_1

- qt=5.9.7=h5867ecd_1

- readline=8.0=h7b6447c_0

- requests=2.28.1=py37h06a4308_0

- scikit-learn=0.23.1=py37h423224d_0

- scipy=1.5.0=py37h0b6359f_0

- setuptools=47.3.1=py37_0

- sip=4.19.8=py37hf484d3e_0

- six=1.15.0=py_0

- snappy=1.1.8=he6710b0_0

- sqlite=3.32.3=h62c20be_0

- threadpoolctl=2.1.0=pyh5ca1d4c_0

- tk=8.6.10=hbc83047_0

- torchvision=0.14.1=py37_cpu

- tornado=6.0.4=py37h7b6447c_1

- urllib3=1.26.14=py37h06a4308_0

- wheel=0.34.2=py37_0

- xz=5.2.5=h7b6447c_0

- yaml=0.2.5=h7b6447c_0

- zlib=1.2.11=h7b6447c_3

- zstd=1.4.4=h0b5b093_3

- pip:

- decorator==5.1.1

- jinja2==3.1.4

- markupsafe==2.1.5

- mne==1.3.1

- packaging==24.0

- platformdirs==4.0.0

- pooch==1.8.2

- pyedflib==0.1.37

- tqdm==4.66.4

- typing-extensions==4.7.1

prefix: /zhome/dd/4/109414/miniconda3/envs/grieger
