## Supplementary material for "Lack of Consensus for Manual Mouse Sleep Scoring Limits Implementation of Automatic Deep Learning Models": environment_slumber

**Data S3: Environment.txt file for SlumberNet model**

name: slumbernet

channels:

- conda-forge

- defaults

dependencies:

- _libgcc_mutex=0.1=conda_forge

- _openmp_mutex=4.5=2_gnu

- _tflow_select=2.3.0=mkl

- abseil-cpp=20211102.0=hd4dd3e8_0

- absl-py=2.1.0=py310h06a4308_0

- aiohttp=3.9.5=py310h5eee18b_0

- aiosignal=1.2.0=pyhd3eb1b0_0

- anyio=4.2.0=py310h06a4308_0

- aom=3.5.0=h27087fc_0

- appdirs=1.4.4=pyhd3eb1b0_0

- argon2-cffi=21.3.0=pyhd3eb1b0_0

- argon2-cffi-bindings=21.2.0=py310h7f8727e_0

- asttokens=2.0.5=pyhd3eb1b0_0

- astunparse=1.6.3=py_0

- async-lru=2.0.4=py310h06a4308_0

- async-timeout=4.0.3=py310h06a4308_0

- attrs=23.1.0=py310h06a4308_0

- babel=2.11.0=py310h06a4308_0

- beautifulsoup4=4.12.3=py310h06a4308_0

- blas=1.0=mkl

- bleach=4.1.0=pyhd3eb1b0_0

- blinker=1.6.2=py310h06a4308_0

- blosc=1.21.3=h6a678d5_0

- bottleneck=1.3.7=py310ha9d4c09_0

- brotli=1.0.9=h5eee18b_8

- brotli-bin=1.0.9=h5eee18b_8

- brotli-python=1.0.9=py310h6a678d5_8

- bzip2=1.0.8=h5eee18b_6

- c-ares=1.19.1=h5eee18b_0

- ca-certificates=2024.7.4=hbcca054_0

- cached-property=1.5.2=py_0

- cachetools=5.3.3=py310h06a4308_0

- certifi=2024.7.4=py310h06a4308_0

- cffi=1.16.0=py310h5eee18b_1

- charset-normalizer=3.3.2=pyhd3eb1b0_0

- click=8.1.7=py310h06a4308_0

- cloudpickle=3.0.0=py310h06a4308_0

- comm=0.2.1=py310h06a4308_0

- contourpy=1.2.0=py310hdb19cb5_0

- cryptography=41.0.3=py310h130f0dd_0

- curl=8.2.1=h37d81fd_0

- cycler=0.11.0=pyhd3eb1b0_0

- cyrus-sasl=2.1.28=h9c0eb46_1

- cytoolz=0.12.2=py310h5eee18b_0

- darkdetect=0.8.0=pyhd8ed1ab_0

- dask-core=2024.5.0=py310h06a4308_0

- dbus=1.13.18=hb2f20db_0

- debugpy=1.6.7=py310h6a678d5_0

- decorator=5.1.1=pyhd3eb1b0_0

- deepdiff=7.0.1=py310h2f386ee_0

- defusedxml=0.7.1=pyhd3eb1b0_0

- deprecated=1.2.13=py310h06a4308_0

- dipy=1.9.0=py310hcc13569_0

- docker-pycreds=0.4.0=pyhd3eb1b0_0

- double-conversion=3.2.0=h27087fc_1

- edfio=0.4.3=pyhd8ed1ab_0

- eeglabio=0.0.2.post4=pyhd8ed1ab_0

- eigen=3.4.0=hdb19cb5_0

- elfutils=0.189=hde5d1a3_0

- exceptiongroup=1.2.0=py310h06a4308_0

- executing=0.8.3=pyhd3eb1b0_0

- expat=2.6.2=h6a678d5_0

- ffmpeg=4.4.2=gpl_hbd009f3_109

- flatbuffers=2.0.0=h2531618_0

- font-ttf-dejavu-sans-mono=2.37=hd3eb1b0_0

- font-ttf-inconsolata=2.001=hcb22688_0

- font-ttf-source-code-pro=2.030=hd3eb1b0_0

- font-ttf-ubuntu=0.83=h8b1ccd4_0

- fontconfig=2.14.1=hef1e5e3_0

- fonts-anaconda=1=h8fa9717_0

- fonts-conda-ecosystem=1=hd3eb1b0_0

- fonttools=4.51.0=py310h5eee18b_0

- freetype=2.12.1=h4a9f257_0

- frozenlist=1.4.0=py310h5eee18b_0

- fsspec=2024.3.1=py310h06a4308_0

- future=0.18.3=py310h06a4308_0

- gast=0.4.0=pyhd3eb1b0_0

- giflib=5.2.1=h5eee18b_3

- gitdb=4.0.7=pyhd3eb1b0_0

- gitpython=3.1.43=py310h06a4308_0

- gl2ps=1.4.2=h70c0345_1

- glib=2.78.4=h6a678d5_0

- glib-tools=2.78.4=h6a678d5_0

- gmp=6.2.1=h295c915_3

- gnutls=3.7.9=hb077bed_0

- google-auth=2.29.0=py310h06a4308_0

- google-auth-oauthlib=0.4.4=pyhd3eb1b0_0

- google-pasta=0.2.0=pyhd3eb1b0_0

- grpc-cpp=1.46.1=h33aed49_1

- grpcio=1.42.0=py310hce63b2e_0

- gst-plugins-base=1.14.1=h6a678d5_1

- gstreamer=1.14.1=h5eee18b_1

- h5io=0.2.4=pyhecae5ae_0

- h5py=3.8.0=nompi_py310h0311031_100

- hdf4=4.2.15=h9772cbc_5

- hdf5=1.12.2=nompi_h2386368_101

- icu=58.2=he6710b0_3

- idna=3.7=py310h06a4308_0

- imagecodecs-lite=2019.12.3=py310h261611a_8

- imageio=2.33.1=py310h06a4308_0

- imageio-ffmpeg=0.5.1=pyhd8ed1ab_0

- importlib-metadata=7.0.1=py310h06a4308_0

- importlib_resources=6.4.0=py310h06a4308_0

- intel-openmp=2023.1.0=hdb19cb5_46306

- ipykernel=6.28.0=py310h06a4308_0

- ipython=8.25.0=py310h06a4308_0

- ipywidgets=8.1.2=py310h06a4308_0

- jedi=0.19.1=py310h06a4308_0

- jinja2=3.1.4=py310h06a4308_0

- joblib=1.4.2=pyhd8ed1ab_0

- jpeg=9e=h5eee18b_2

- json5=0.9.6=pyhd3eb1b0_0

- jsoncpp=1.9.5=h4bd325d_1

- jsonschema=4.19.2=py310h06a4308_0

- jsonschema-specifications=2023.7.1=py310h06a4308_0

- jupyter=1.0.0=py310h06a4308_9

- jupyter-lsp=2.2.0=py310h06a4308_0

- jupyter_client=8.6.0=py310h06a4308_0

- jupyter_console=6.6.3=py310h06a4308_0

- jupyter_core=5.7.2=py310h06a4308_0

- jupyter_events=0.10.0=py310h06a4308_0

- jupyter_server=2.14.1=py310h06a4308_0

- jupyter_server_terminals=0.4.4=py310h06a4308_1

- jupyterlab=4.0.11=py310h06a4308_0

- jupyterlab_pygments=0.1.2=py_0

- jupyterlab_server=2.25.1=py310h06a4308_0

- jupyterlab_widgets=3.0.10=py310h06a4308_0

- keras=2.11.0=pyhd8ed1ab_0

- keras-preprocessing=1.1.2=pyhd3eb1b0_0

- kiwisolver=1.4.4=py310h6a678d5_0

- krb5=1.20.1=h568e23c_1

- lame=3.100=h7b6447c_0

- lazy_loader=0.4=py310h06a4308_0

- lcms2=2.12=h3be6417_0

- ld_impl_linux-64=2.38=h1181459_1

- lerc=3.0=h295c915_0

- libaec=1.1.3=h59595ed_0

- libarchive=3.6.2=hab531cd_0

- libblas=3.9.0=1_h86c2bf4_netlib

- libbrotlicommon=1.0.9=h5eee18b_8

- libbrotlidec=1.0.9=h5eee18b_8

- libbrotlienc=1.0.9=h5eee18b_8

- libcblas=3.9.0=6_ha36c22a_netlib

- libclang=14.0.6=default_hc6dbbc7_1

- libclang13=14.0.6=default_he11475f_1

- libcups=2.4.2=ha637b67_0

- libcurl=8.2.1=h91b91d3_0

- libdeflate=1.8=h7f8727e_5

- libdrm=2.4.122=h4ab18f5_0

- libedit=3.1.20230828=h5eee18b_0

- libev=4.33=h7f8727e_1

- libexpat=2.6.2=h59595ed_0

- libffi=3.4.4=h6a678d5_1

- libgcc-ng=14.1.0=h77fa898_0

- libgfortran=3.0.0=1

- libgfortran-ng=14.1.0=h69a702a_0

- libgfortran5=14.1.0=hc5f4f2c_0

- libglib=2.78.4=hdc74915_0

- libgomp=14.1.0=h77fa898_0

- libiconv=1.16=h5eee18b_3

- libidn2=2.3.4=h5eee18b_0

- liblapack=3.9.0=6_ha36c22a_netlib

- libllvm14=14.0.6=hdb19cb5_3

- libllvm17=17.0.6=hc9c083f_0

- libmatio=1.5.23=h63e3022_1

- libmicrohttpd=0.9.77=h97afed2_0

- libnetcdf=4.8.1=nompi_h21705cb_104

- libnghttp2=1.52.0=ha637b67_1

- libnsl=2.0.0=h5eee18b_0

- libogg=1.3.5=h27cfd23_1

- libopenblas=0.3.21=h043d6bf_0

- libpciaccess=0.18=hd590300_0

- libpng=1.6.39=h5eee18b_0

- libpq=12.15=h37d81fd_1

- libprotobuf=3.20.3=he621ea3_0

- libsodium=1.0.18=h7b6447c_0

- libsqlite=3.46.0=hde9e2c9_0

- libssh2=1.10.0=h37d81fd_2

- libstdcxx-ng=14.1.0=hc0a3c3a_0

- libtasn1=4.19.0=h5eee18b_0

- libtheora=1.1.1=h7f8727e_3

- libtiff=4.4.0=hecacb30_2

- libunistring=0.9.10=h27cfd23_0

- libuuid=2.38.1=h0b41bf4_0

- libva=2.21.0=h4ab18f5_2

- libvorbis=1.3.7=h7b6447c_0

- libvpx=1.11.0=h295c915_0

- libwebp-base=1.3.2=h5eee18b_0

- libxcb=1.15=h7f8727e_0

- libxkbcommon=1.0.1=hfa300c1_0

- libxml2=2.9.14=h74e7548_0

- libxslt=1.1.35=h4e12654_0

- libzip=1.9.2=hc869a4a_1

- libzlib=1.2.13=h4ab18f5_6

- llvmlite=0.43.0=py310h6a678d5_0

- locket=1.0.0=py310h06a4308_0

- loguru=0.5.3=py310h06a4308_4

- lxml=4.9.1=py310h1edc446_0

- lz4-c=1.9.4=h6a678d5_1

- lzo=2.10=h7b6447c_2

- markdown=3.4.1=py310h06a4308_0

- markupsafe=2.1.3=py310h5eee18b_0

- matplotlib=3.8.4=py310h06a4308_0

- matplotlib-base=3.8.4=py310h1128e8f_0

- matplotlib-inline=0.1.6=py310h06a4308_0

- mesalib=23.3.2=h6b56f8e_0

- mffpy=0.9.0=pyhd8ed1ab_0

- mistune=2.0.4=py310h06a4308_0

- mkl=2023.1.0=h213fc3f_46344

- mkl-service=2.4.0=py310h5eee18b_1

- mkl_fft=1.3.8=py310h5eee18b_0

- mkl_random=1.2.4=py310hdb19cb5_0

- mne=1.7.1=pyqt_h499dab4_202

- mne-base=1.7.1=pyha770c72_202

- mne-qt-browser=0.6.3=pyha770c72_0

- more-itertools=10.1.0=py310h06a4308_0

- msgpack-python=1.0.3=py310hd09550d_0

- multidict=6.0.4=py310h5eee18b_0

- mysql=5.7.24=he378463_2

- nbclient=0.8.0=py310h06a4308_0

- nbconvert=7.10.0=py310h06a4308_0

- nbformat=5.9.2=py310h06a4308_0

- ncurses=6.4=h6a678d5_0

- nest-asyncio=1.6.0=py310h06a4308_0

- nettle=3.9.1=h7ab15ed_0

- networkx=3.3=py310h06a4308_0

- nibabel=5.2.1=pyha770c72_0

- nilearn=0.10.4=pyhd8ed1ab_0

- nlohmann_json=3.11.2=h6a678d5_0

- notebook=7.0.8=py310h06a4308_2

- notebook-shim=0.2.3=py310h06a4308_0

- numba=0.60.0=py310h5dc88bb_0

- numexpr=2.8.7=py310h85018f9_0

- numpy=1.26.4=py310h5f9d8c6_0

- numpy-base=1.26.4=py310hb5e798b_0

- oauthlib=3.2.2=py310h06a4308_0

- openh264=2.3.1=hcb278e6_2

- openjpeg=2.4.0=h9ca470c_2

- openmeeg=2.5.6=py310h31ff90e_0

- openssl=1.1.1w=h7f8727e_0

- opt_einsum=3.3.0=pyhd3eb1b0_1

- ordered-set=4.1.0=py310h06a4308_0

- orjson=3.9.15=py310h97a8848_0

- overrides=7.4.0=py310h06a4308_0

- p11-kit=0.24.1=hc5aa10d_0

- packaging=24.1=py310h06a4308_0

- pandas=2.1.4=py310h1128e8f_0

- pandocfilters=1.5.0=pyhd3eb1b0_0

- parso=0.8.3=pyhd3eb1b0_0

- partd=1.4.1=py310h06a4308_0

- pathtools=0.1.2=pyhd3eb1b0_1

- patsy=0.5.6=py310h06a4308_0

- pcre2=10.42=hebb0a14_1

- pexpect=4.8.0=pyhd3eb1b0_3

- pillow=10.4.0=py310h5eee18b_0

- pip=24.0=py310h06a4308_0

- platformdirs=3.10.0=py310h06a4308_0

- ply=3.11=py310h06a4308_0

- pooch=1.7.0=py310h06a4308_0

- proj=9.1.0=h93bde94_0

- prometheus_client=0.14.1=py310h06a4308_0

- prompt-toolkit=3.0.43=py310h06a4308_0

- prompt_toolkit=3.0.43=hd3eb1b0_0

- protobuf=3.20.3=py310h6a678d5_0

- psutil=5.9.0=py310h5eee18b_0

- ptyprocess=0.7.0=pyhd3eb1b0_2

- pugixml=1.11.4=h295c915_1

- pure_eval=0.2.2=pyhd3eb1b0_0

- pyasn1=0.4.8=pyhd3eb1b0_0

- pyasn1-modules=0.2.8=py_0

- pybv=0.7.5=pyhd8ed1ab_0

- pycparser=2.21=pyhd3eb1b0_0

- pygments=2.15.1=py310h06a4308_1

- pyjwt=2.8.0=py310h06a4308_0

- pymatreader=0.0.32=pyhd8ed1ab_0

- pyopengl=3.1.1a1=py310h06a4308_0

- pyopenssl=23.2.0=py310h06a4308_0

- pyparsing=3.0.9=py310h06a4308_0

- pyqt=5.15.10=py310h6a678d5_0

- pyqt5-sip=12.13.0=py310h5eee18b_0

- pyqtgraph=0.13.1=py310h06a4308_0

- pysocks=1.7.1=py310h06a4308_0

- pytables=3.7.0=py310hb60b9b2_3

- python=3.10.8=h257c98d_0_cpython

- python-dateutil=2.9.0post0=py310h06a4308_2

- python-fastjsonschema=2.16.2=py310h06a4308_0

- python-flatbuffers=2.0=pyhd3eb1b0_0

- python-json-logger=2.0.7=py310h06a4308_0

- python-picard=0.7=pyh8a188c0_0

- python-tzdata=2023.3=pyhd3eb1b0_0

- python_abi=3.10=2_cp310

- pytz=2024.1=py310h06a4308_0

- pyvista=0.44.1=pyhd8ed1ab_0

- pyvistaqt=0.11.1=pyhd8ed1ab_0

- pywavelets=1.5.0=py310ha9d4c09_0

- pyyaml=6.0.1=py310h5eee18b_0

- pyzmq=25.1.2=py310h6a678d5_0

- qdarkstyle=3.2.3=pyhd3eb1b0_0

- qt-main=5.15.2=h5b8104b_9

- qtconsole=5.5.1=py310h06a4308_0

- qtpy=2.4.1=py310h06a4308_0

- re2=2022.04.01=h295c915_0

- readline=8.2=h5eee18b_0

- referencing=0.30.2=py310h06a4308_0

- requests=2.32.3=py310h06a4308_0

- requests-oauthlib=2.0.0=py310h06a4308_0

- rfc3339-validator=0.1.4=py310h06a4308_0

- rfc3986-validator=0.1.1=py310h06a4308_0

- rpds-py=0.10.6=py310hb02cf49_0

- rsa=4.7.2=pyhd3eb1b0_1

- scikit-image=0.20.0=py310h6a678d5_0

- scikit-learn=1.5.1=py310h146d792_0

- scipy=1.14.0=py310h93e2701_1

- scooby=0.10.0=pyhd8ed1ab_0

- seaborn=0.13.2=py310h06a4308_0

- send2trash=1.8.2=py310h06a4308_0

- sentry-sdk=1.9.0=py310h06a4308_0

- setproctitle=1.2.2=py310h7f8727e_0

- setuptools=69.5.1=py310h06a4308_0

- setuptools-scm=8.1.0=py310h06a4308_0

- sip=6.7.12=py310h6a678d5_0

- six=1.16.0=pyhd3eb1b0_1

- smmap=4.0.0=pyhd3eb1b0_0

- snappy=1.1.10=h6a678d5_1

- sniffio=1.3.0=py310h06a4308_0

- soupsieve=2.5=py310h06a4308_0

- sqlite=3.45.3=h5eee18b_0

- stack_data=0.2.0=pyhd3eb1b0_0

- statsmodels=0.14.2=py310h5eee18b_0

- svt-av1=1.3.0=h27087fc_0

- tbb=2021.8.0=hdb19cb5_0

- tbb-devel=2021.8.0=hdb19cb5_0

- tensorboard=2.11.0=py310h06a4308_0

- tensorboard-data-server=0.6.1=py310h52d8a92_0

- tensorboard-plugin-wit=1.8.1=py310h06a4308_0

- tensorflow=2.11.0=mkl_py310hb40ee82_0

- tensorflow-base=2.11.0=mkl_py310he5f8e37_0

- tensorflow-estimator=2.11.0=py310h06a4308_0

- termcolor=2.1.0=py310h06a4308_0

- terminado=0.17.1=py310h06a4308_0

- threadpoolctl=3.5.0=pyhc1e730c_0

- tifffile=2020.6.3=py_0

- tinycss2=1.2.1=py310h06a4308_0

- tk=8.6.14=h39e8969_0

- tomli=2.0.1=py310h06a4308_0

- toolz=0.12.0=py310h06a4308_0

- tornado=6.4.1=py310h5eee18b_0

- tqdm=4.66.4=py310h2f386ee_0

- traitlets=5.14.3=py310h06a4308_0

- trame=3.6.3=pyhd8ed1ab_0

- trame-client=3.2.1=pyhd8ed1ab_0

- trame-server=3.0.3=pyhd8ed1ab_0

- trame-vtk=2.8.9=pyhd8ed1ab_0

- trame-vuetify=2.6.2=pyhd8ed1ab_0

- trx-python=0.3=py310hff52083_0

- typing-extensions=4.11.0=py310h06a4308_0

- typing_extensions=4.11.0=py310h06a4308_0

- tzdata=2024a=h04d1e81_0

- unicodedata2=15.1.0=py310h5eee18b_0

- urllib3=2.2.2=py310h06a4308_0

- utfcpp=3.2.1=h06a4308_0

- vtk=9.2.2=osmesa_py310h47209f7_102

- wandb=0.16.6=pyhd8ed1ab_0

- wcwidth=0.2.5=pyhd3eb1b0_0

- webencodings=0.5.1=py310h06a4308_1

- websocket-client=1.8.0=py310h06a4308_0

- werkzeug=3.0.3=py310h06a4308_0

- wheel=0.43.0=py310h06a4308_0

- widgetsnbextension=4.0.10=py310h06a4308_0

- wrapt=1.14.1=py310h5eee18b_0

- wslink=2.1.1=pyhd8ed1ab_0

- x264=1!164.3095=h166bdaf_2

- x265=3.5=h924138e_3

- xlrd=2.0.1=pyhd3eb1b0_1

- xmltodict=0.13.0=py310h06a4308_0

- xorg-damageproto=1.2.1=h7f98852_1002

- xorg-fixesproto=5.0=h7f98852_1002

- xorg-glproto=1.4.17=h7f98852_1002

- xorg-kbproto=1.0.7=h7f98852_1002

- xorg-libice=1.1.1=hd590300_0

- xorg-libsm=1.2.4=h7391055_0

- xorg-libx11=1.8.9=h8ee46fc_0

- xorg-libxdamage=1.1.5=h7f98852_1

- xorg-libxext=1.3.4=h0b41bf4_2

- xorg-libxfixes=5.0.3=h7f98852_1004

- xorg-libxrandr=1.5.2=h7f98852_1

- xorg-libxrender=0.9.11=hd590300_0

- xorg-libxt=1.3.0=hd590300_1

- xorg-randrproto=1.5.0=h7f98852_1001

- xorg-renderproto=0.11.1=h7f98852_1002

- xorg-util-macros=1.19.0=h27cfd23_2

- xorg-xextproto=7.3.0=h0b41bf4_1003

- xorg-xf86vidmodeproto=2.3.1=h7f98852_1002

- xorg-xproto=7.0.31=h27cfd23_1007

- xz=5.4.6=h5eee18b_1

- yaml=0.2.5=h7b6447c_0

- yarl=1.9.3=py310h5eee18b_0

- zeromq=4.3.5=h6a678d5_0

- zipp=3.17.0=py310h06a4308_0

- zlib=1.2.13=h4ab18f5_6

- zstd=1.5.5=hc292b87_2

prefix: /zhome/dd/4/109414/miniconda3/envs/slumbernet
