## Supplementary material for "Lack of Consensus for Manual Mouse Sleep Scoring Limits Implementation of Automatic Deep Learning Models": environment_spindle

**Data S1: Environment.txt file for SPINDLE model**

name: spindle_javier

channels:

- anaconda

- defaults

dependencies:

- _libgcc_mutex=0.1=main

- _openmp_mutex=5.1=1_gnu

- blas=1.0=mkl

- blosc=1.21.3=h6a678d5_0

- bottleneck=1.3.5=py39h7deecbd_0

- brotli=1.0.9=h5eee18b_8

- brotli-bin=1.0.9=h5eee18b_8

- bzip2=1.0.8=h5eee18b_6

- c-ares=1.19.1=h5eee18b_0

- c-blosc2=2.12.0=h80c7b02_0

- ca-certificates=2024.9.24=h06a4308_0

- contourpy=1.2.0=py39hdb19cb5_0

- freetype=2.12.1=h4a9f257_0

- hdf5=1.12.1=h2b7332f_3

- importlib_resources=6.1.1=py39h06a4308_1

- intel-openmp=2023.1.0=hdb19cb5_46306

- joblib=1.2.0=py39h06a4308_0

- jpeg=9e=h5eee18b_1

- krb5=1.20.1=h143b758_1

- lcms2=2.12=h3be6417_0

- ld_impl_linux-64=2.38=h1181459_1

- lerc=3.0=h295c915_0

- libbrotlicommon=1.0.9=h5eee18b_8

- libbrotlidec=1.0.9=h5eee18b_8

- libbrotlienc=1.0.9=h5eee18b_8

- libcurl=8.7.1=h251f7ec_0

- libdeflate=1.17=h5eee18b_1

- libedit=3.1.20230828=h5eee18b_0

- libev=4.33=h7f8727e_1

- libffi=3.4.4=h6a678d5_0

- libgcc-ng=11.2.0=h1234567_1

- libgfortran-ng=11.2.0=h00389a5_1

- libgfortran5=11.2.0=h1234567_1

- libgomp=11.2.0=h1234567_1

- libnghttp2=1.57.0=h2d74bed_0

- libpng=1.6.39=h5eee18b_0

- libssh2=1.11.0=h251f7ec_0

- libstdcxx-ng=11.2.0=h1234567_1

- libtiff=4.5.1=h6a678d5_0

- libwebp-base=1.3.2=h5eee18b_0

- lz4-c=1.9.4=h6a678d5_1

- lzo=2.10=h7b6447c_2

- matplotlib-base=3.8.4=py39h1128e8f_0

- mkl=2023.1.0=h213fc3f_46344

- mkl-service=2.4.0=py39h5eee18b_1

- mkl_fft=1.3.8=py39h5eee18b_0

- mkl_random=1.2.4=py39hdb19cb5_0

- ncurses=6.4=h6a678d5_0

- numexpr=2.8.7=py39h85018f9_0

- numpy=1.26.2=py39h5f9d8c6_0

- numpy-base=1.26.2=py39hb5e798b_0

- openjpeg=2.4.0=h3ad879b_0

- openssl=3.0.15=h5eee18b_0

- packaging=23.2=py39h06a4308_0

- pandas=2.1.4=py39h1128e8f_0

- patsy=0.5.6=py39h06a4308_0

- pip=23.3.1=py39h06a4308_0

- py-cpuinfo=9.0.0=py39h06a4308_0

- python=3.9.18=h955ad1f_0

- python-dateutil=2.8.2=pyhd3eb1b0_0

- python-tzdata=2023.3=pyhd3eb1b0_0

- pytz=2023.3.post1=py39h06a4308_0

- pyyaml=6.0.1=py39h5eee18b_0

- readline=8.2=h5eee18b_0

- scikit-learn=1.3.0=py39h1128e8f_0

- scipy=1.11.4=py39h5f9d8c6_0

- seaborn=0.13.2=py39h06a4308_0

- setuptools=68.2.2=py39h06a4308_0

- six=1.16.0=pyhd3eb1b0_1

- sqlite=3.41.2=h5eee18b_0

- statsmodels=0.14.2=py39h5eee18b_0

- tables=3.9.2=py39h0016290_0

- tbb=2021.8.0=hdb19cb5_0

- threadpoolctl=2.2.0=pyh0d69192_0

- tk=8.6.12=h1ccaba5_0

- tzdata=2023c=h04d1e81_0

- unicodedata2=15.1.0=py39h5eee18b_0

- wheel=0.41.2=py39h06a4308_0

- xz=5.4.5=h5eee18b_0

- yaml=0.2.5=h7b6447c_0

- zipp=3.17.0=py39h06a4308_0

- zlib=1.2.13=h5eee18b_0

- zlib-ng=2.0.7=h5eee18b_0

- zstd=1.5.5=hc292b87_0

- pip:

- absl-py==2.0.0

- astunparse==1.6.3

- cachetools==5.3.2

- certifi==2023.11.17

- charset-normalizer==3.3.2

- click==8.1.7

- cycler==0.12.1

- decorator==5.1.1

- defusedxml==0.7.1

- docker-pycreds==0.4.0

- et-xmlfile==1.1.0

- flatbuffers==23.5.26

- fonttools==4.47.0

- gast==0.5.4

- gitdb==4.0.11

- gitpython==3.1.43

- google-auth==2.25.2

- google-auth-oauthlib==1.2.0

- google-pasta==0.2.0

- grpcio==1.60.0

- h5py==3.10.0

- idna==3.6

- importlib-metadata==7.0.1

- jinja2==3.1.2

- keras==2.15.0

- kiwisolver==1.4.5

- lazy-loader==0.3

- libclang==16.0.6

- markdown==3.5.1

- markupsafe==2.1.3

- matplotlib==3.8.2

- ml-dtypes==0.2.0

- mne==1.6.0

- nvidia-cublas-cu12==12.2.5.6

- nvidia-cuda-cupti-cu12==12.2.142

- nvidia-cuda-nvcc-cu12==12.2.140

- nvidia-cuda-nvrtc-cu12==12.2.140

- nvidia-cuda-runtime-cu12==12.2.140

- nvidia-cudnn-cu12==8.9.4.25

- nvidia-cufft-cu12==11.0.8.103

- nvidia-curand-cu12==10.3.3.141

- nvidia-cusolver-cu12==11.5.2.141

- nvidia-cusparse-cu12==12.1.2.141

- nvidia-nccl-cu12==2.16.5

- nvidia-nvjitlink-cu12==12.2.140

- oauthlib==3.2.2

- openpyxl==3.1.5

- opt-einsum==3.3.0

- pdf2image==1.17.0

- pillow==10.1.0

- platformdirs==4.1.0

- pooch==1.8.0

- protobuf==4.23.4

- psutil==6.0.0

- pyasn1==0.5.1

- pyasn1-modules==0.3.0

- pyparsing==3.1.1

- requests==2.31.0

- requests-oauthlib==1.3.1

- rsa==4.9

- sentry-sdk==2.7.1

- setproctitle==1.3.3

- smmap==5.0.1

- tensorboard==2.15.1

- tensorboard-data-server==0.7.2

- tensorflow==2.15.0.post1

- tensorflow-estimator==2.15.0

- tensorflow-io-gcs-filesystem==0.35.0

- termcolor==2.4.0

- torchsummary==1.5.1

- tqdm==4.66.1

- typing-extensions==4.9.0

- urllib3==2.1.0

- wandb==0.17.3

- werkzeug==3.0.1

- wrapt==1.14.1

prefix: /zhome/dd/4/109414/miniconda3/envs/spindle_javier
